# Inflammasomes and DNA Damage Orchestrate Divergent Pyroptotic Fates in Neutrophils

**DOI:** 10.1101/2025.09.03.673694

**Authors:** Apurwa Singhal, Neha Awasthi, Partha Sarathi Mondal, Nishakumari Chentunarayan Singh, Ritvik Guru, Tulika Chandra, Kalyan Mitra, Madhu Dikshit, Sachin Kumar

## Abstract

The understanding of pyroptotic cell death and its mechanistic insights in neutrophils remains less explored and ambiguous. This study analyzes neutrophil pyroptotic responses under different conditions and dissects distinct pyroptosis-associated events, including IL-1β release and cell death. We decipher the signalling pathways underlying these responses and distinguish them from other cell death mechanisms. Interestingly, we observe IL-1β release and enhanced survival in neutrophils in response to LPS-primed ATP-treatment, in contrast to a combination of IL-1β release with cell death in macrophages. While LPS-primed nigericin-treated neutrophils exhibit IL-1β release and cell death, characterized by nuclear rounding, cell swelling/ ballooning, plasma membrane pore formation, and subsequent cell rupture, confirming the occurrence of pyroptotic cell death. While nigericin alone triggers cytokine uncoupled pyroptosis. Intriguingly, these phenomena do not induce under other death programs, including apoptosis, NETosis. This provides opportunity to unravel the regulation of these pyroptotic responses in neutrophils. Data observed reveal the role of NLRP3 and context-dependent caspase −1 & 11 in IL-1β secretion. While, DNA damage and caspase-7 & 9 regulate pyroptotic death. Intriguingly, forced DNA damage by ATM kinase inhibitor mitigated inflammasome activation and IL-1β release, while spur death. Furthermore, this study identifies perinuclear actomyosin forces driving nuclear rounding and pyroptotic death. Both murine neutrophils exposed to bacteria and human neutrophils treated with nigericin exhibit these pyroptotic processes, highlighting their broad relevance. This study depicts decision whether to secrete cytokines or undergo pyroptotic cell death emphasizing the regulatory role of DNA damage-cytoskeletal axis beyond inflammasome activation. Moreover, LPS and bacteria induced acute lung injury in mice displays nuclear rounding presenting pyroptotic neutrophils and enhanced NLRP3, Caspase-11 activation, and IL-1β release. Together, this study defines intriguing crosstalk of NLRP3, caspases, DNA damage, and actomyosin forces that orchestrate divergent inflammatory fates, potentially leading to opportunities for targeted strategies for combating inflammation.

**HIGHLIGHTS:**

- Neutrophils undergo pyroptotic cell death with nigericin and bacteria, but not with ATP.
- LPS priming is essential for IL-1β secretion, while dispensable for cell swelling/ballooning and death.
- NLRP3 and Caspase 1/11 regulate IL-1β secretion, while DNA damage and Caspase-7/9 control pyroptotic cell death.
- Perinuclear actomyosin signalling regulates nuclear rounding during pyroptotic cell death.

**GRAPHICAL ABSTRACT:** 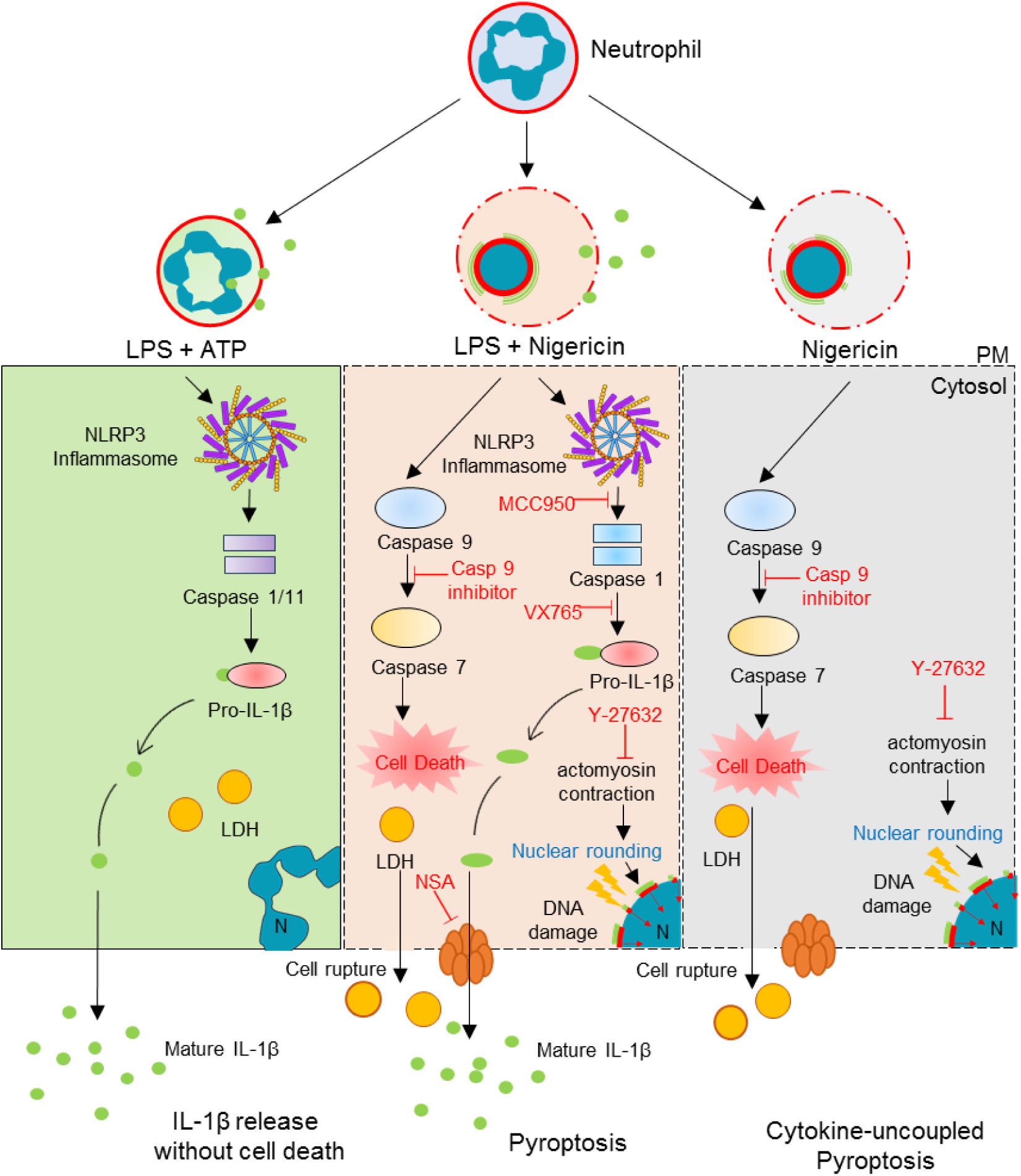

## INTRODUCTION

Polymorphonuclear neutrophils (PMNs) are the key immune cells that sense and fight infections caused by bacterial, fungal, and other pathogens. These cells eliminate intruders by phagocytosis, and neutrophil extracellular traps (NETs) (1, 2). Neutrophils release oxidants, toxic proteases, and antigenic chromatin via over-activation and distinct forms of death that can also drive various inflammatory diseases, including vasculitis, systemic lupus erythematosus, COVID-19 and sepsis (2–4). Thus, understanding distinct neutrophil deaths and the mechanistic insights is warranted for therapeutic purposes. These short-lived cells often die through non-lytic, anti-inflammatory apoptosis under homeostatic conditions, while under different inflammatory conditions can undergo lytic death programs, including NETosis and necroptosis, associated with inflammatory outcomes (2, 4). Another lytic cell program, pyroptosis, is a highly inflammatory death, averting the intracellular growth of pathogens (5). The understanding of pyroptosis is primarily driven by macrophages, which remains context-dependent and multifaceted in neutrophils (6–8).

The Greek term pyroptosis reflects ‘pyro’ = fire/ fever, and ‘ptosis’= dropping/falling (9). Pyroptosis is often categorized by the release of pro-inflammatory cytokines, including Interleukin-1 beta (IL-1β), Interleukin-18 (IL-18), and cell swelling/ ballooning, followed by rupture (9). Mechanistically, pathogen recognition receptors lead to pro-IL-1β transcription via a multi-protein Nod-like receptor (NLR) family pyrin domain-containing 3 (NLRP3) inflammasome complex. Furthermore, activated caspases −1, 4, 6 & 11 process pro-IL-1β into mature IL-1β. These caspases cleave Gasdermin D (GSDMD) to N-GSDMD subunits, leading to plasma membrane pore formation, IL-1β release, and pyroptotic cell death (5, 10–12). However, recent studies suggest that cell lysis and IL-1β secretion can be uncoupled and occur independently. For example, IL-1β secretion does not necessarily require priming in monocytes (13) and is also released from live non-pyroptotic cells, including macrophages and dendritic cells (7, 14–16).

Neutrophils release substantial amounts of IL-1β at sites of infection and inflammation that regulate pathological outcomes (7, 17). Intracellular sensors, such as Nod-like receptors (NLRs), including NLRP3, NLR family CARD domain-containing protein 4 (NLRC4), and absent in melanoma 2 (AIM2), govern IL-1β secretion in murine and human neutrophils (7, 17–19). Moreover, studies showed IL-1β release in the absence of cell death, suggesting neutrophils being resistant to pyroptosis (6, 7, 20). Intriguingly, other studies observed pyroptotic cell death in neutrophils under specific conditions (8, 19, 21, 22). The IL-1β release can also be regulated through Gasdermin-D-dependent and - independent mechanisms (16, 23, 24). Remarkably, N-GSDMD was observed in the membrane of primary granules and autophagosomes in neutrophils, instead of the plasma membrane as shown in other cells, including macrophages (6). This emphasizes the unique regulation of pyroptotic cell death in neutrophils, which is still not well understood. Furthermore, the understanding of pyroptosis, including in neutrophils, is driven by surrogate markers, including IL-1β and lactate dehydrogenase (LDH) release, and activation of NLRPs-caspases-GSDMs signaling in response to various stimuli. While subcellular features and dynamics including cell swelling, pore formation, and membrane rupture, that categorize and distinguish pyroptosis from other forms of cell death, remain often neglected.

In the present study, we investigated and characterized distinct pyroptosis-associated events in murine and human neutrophils through morphological, biochemical, and functional analyses under different conditions. We observed differential responses, precisely referred to “IL-1β release without cell death”, which describes the condition in which IL-1β is released without any cell death. The “pyroptosis” presented a condition that exhibits cell swelling and cell death along with IL-1β release. In contrast, a situation displaying cell swelling and cell death without IL-1β and TNF-α release is described as “cytokine uncoupled pyroptosis”. In this study, we observed that LPS-primed ATP-treated neutrophils release IL-1β and exhibit enhanced survival. In contrast, LPS-primed nigericin-treated neutrophils exhibited pyroptosis with both IL-1β release and pyroptotic cell death. Furthermore, LPS priming was observed to be essential for IL-1β release, but not for cell swelling and pyroptotic cell death in nigericin-treated neutrophils. Interestingly, nigericin triggered cytokine uncoupled pyroptosis. The mechanistic analysis identified NLRP3 and DNA damage as critical determinants of neutrophil pyroptotic signaling outcomes, including IL-1β release and cell death, respectively. Furthermore, this study unravels cytoskeletal dynamics mediated nuclear reorganization, which instigates cell death in response to nigericin and bacteria in neutrophils.

## RESULTS

### Neutrophils exhibit discrete IL-1β release and pyroptotic cell death responses upon ATP and nigericin challenge

To explore the paradox of pyroptosis in neutrophils, we investigated the features associated with distinct pyroptotic processes in murine neutrophils that were untreated or primed with toll-like receptor 4 (TLR-4) ligand, LPS, and subsequently stimulated with inflammasome activators, including nigericin and ATP. Contrary to the previous notions (6, 20), nigericin treatment significantly increased cell death after 3 hours in neutrophils, as assessed by Hoechst-33342 and Sytox Green labelling (**Fig 1A, B)**. Similar results were observed with propidium iodide labelling **(Fig S1A, B)**, and ethidium bromide (EtBr; not shown), demonstrating increased plasma-membrane permeability following nigericin treatment. Conversely, no cell death was observed under conditions involving LPS priming, ATP treatment or LPS-primed ATP treatment (**Fig 1B and Fig S1A, B)**. The Sytox Green kinetics further confirmed a time-dependent increase in death of nigericin-treated neutrophils independent of LPS priming, while no cell death was observed in LPS-primed ATP-treated neutrophils (**Fig 1C**). Interestingly, LPS-primed ATP-treated neutrophils were protected from spontaneous death observed at an extended period of 12 hours (**Fig 1D**). These differential findings observed with nigericin and ATP in LPS-primed neutrophils were further validated through a comparative analysis with macrophages (**Fig 1E**). As expected, at 1 hour time point, LPS-primed macrophages were quite sensitive to both ATP and nigericin treatments, exhibiting over 70% death (6, 16), while neutrophils showed resistance to cell death. An extension of time to 3 hours increased the cell death to 40% in nigericin-treated neutrophils, but ATP-treated neutrophils remained viable (**Fig 1E**), confirming distinct pyroptotic responses in neutrophils compared to macrophages. Additionally, LDH, a determinant of plasma membrane rupture, was significantly increased with nigericin stimulation independent of LPS priming, while no LDH release was observed with LPS+ATP (**Fig 1F and Fig S1C)**. Altogether, neutrophils responded differently to nigericin and ATP stimulation toward the cell death program.

**Fig 1:**
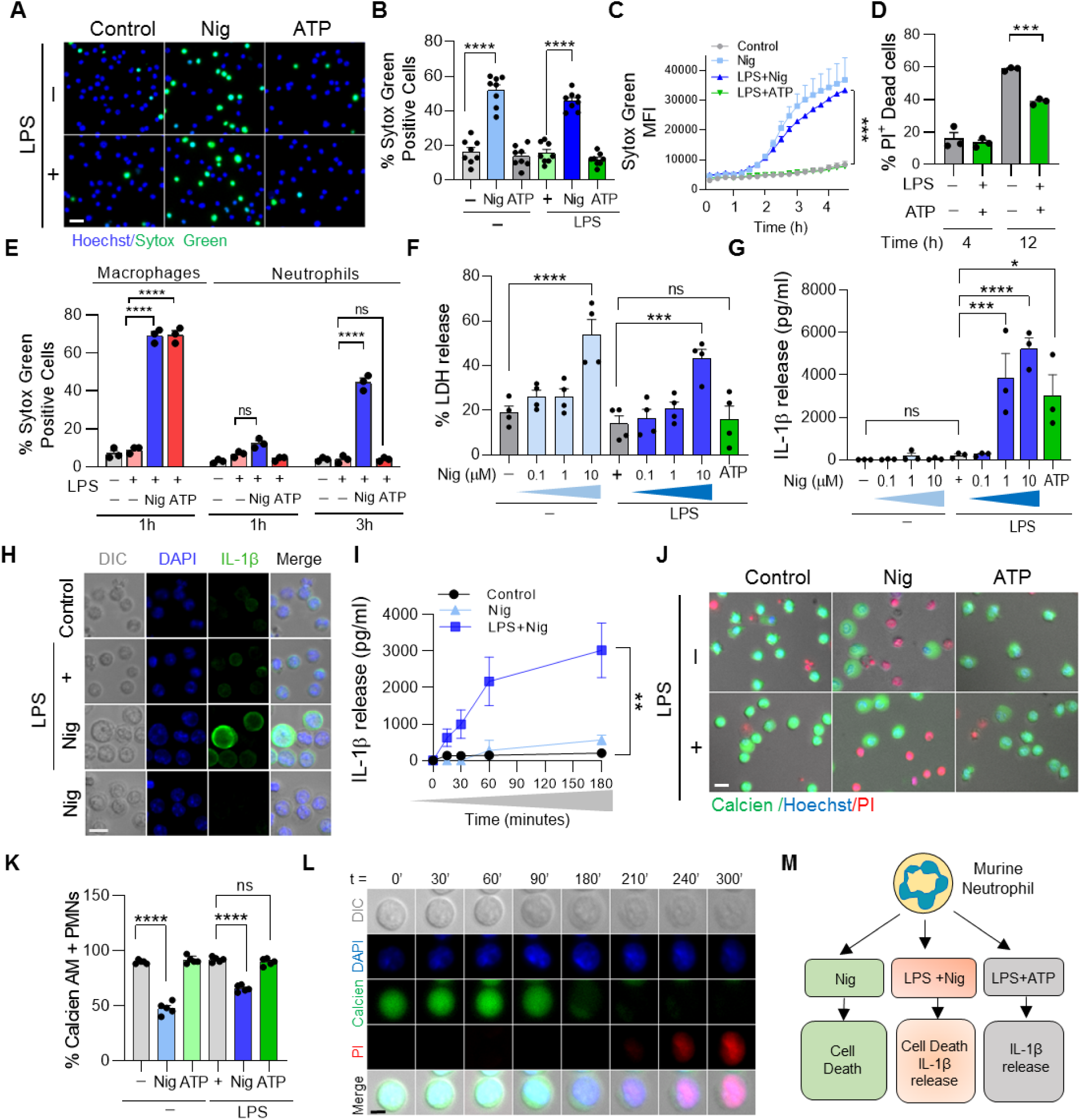
Analysis of cell death and IL-1β responses to Nigericin and ATP in murine neutrophils under with/without LPS-priming settings. (A) Representative image showing Hoechst (blue) and Sytox Green (green) positive cells after 3h of nigericin (10 µM) or ATP (5 mM) treatments in non-primed and LPS-primed neutrophils (Scale bar, 20 μm). (B) Quantification showing Sytox Green positive cells after 3h of nigericin (10 µM) or ATP (5 mM) treatments in non-primed and LPS-primed neutrophils (****P <0.0001, n = 7 independent experiments, analyzed using One-way ANOVA). (C) Time-dependent kinetics for Sytox Green mean fluorescence intensity after nigericin or ATP challenge in non-primed and LPS-primed neutrophils (***P <0.001, n = 3 independent experiments, analyzed using One-way ANOVA). (D) PI uptake in LPS-primed neutrophils after 4 h and 12 h with and without ATP (***P <0.001, n = 3 independent experiments, analyzed using One-way ANOVA). (E) Relative quantification of Sytox Green positive cells in LPS primed neutrophils and peritoneal macrophages after nigericin and ATP treatments (****P <0.0001, ns - not significant, n = 3 independent experiments, analyzed using One-way ANOVA). (F) % LDH release with different doses of nigericin and ATP in non-primed and LPS-primed neutrophils (***P <0.001 and ****P <0.0001, n = 3 independent experiments, analyzed using One-way ANOVA). (G) IL-1β (pg/ml) release with different doses of nigericin or ATP in non-primed and LPS-primed neutrophils (*P<0.05, ***P <0.001, and ****P <0.0001, n = 3 independent experiments, analyzed using One-way ANOVA). (H) Immunofluorescence image showing DIC image (gray), intracellular IL-1β (green), and DAPI (blue) in non-primed and LPS-primed neutrophils treated with nigericin (Scale bar, 10 μm). (I) Time-dependent analysis of IL-1β (pg/ml) release with nigericin in the presence and absence of LPS-priming in neutrophils (**P <0.01, n = 3 independent experiments, analyzed using Two-way ANOVA). (J) Representative image of Calcien AM (green), Hoechst (Blue), and PI (red) staining in non-primed and LPS-primed neutrophils upon nigericin and ATP treatment (Scale bar, 10 μm). (K) Quantification of Calcien AM positive neutrophils after nigericin and ATP treatment with and without LPS priming (****P <0.0001, n = 3 independent experiments, analyzed using One-way ANOVA). (L) Time-dependent analysis for changes in Calcien AM (green), PI (red), and Hoechst (blue) after nigericin addition in PMNs (Scale bar, 2 μm). (M) Representative diagram showing distinct cell death and IL-1β release phenotypes with LPS+Nig, LPS+ATP, and Nig treatments in neutrophils.

Interestingly, the release of pyroptosis-associated cytokine IL-1β was increased upon nigericin-treatment in LPS-primed neutrophils, but not in the absence of priming (**Fig 1G**). Notably, LPS-primed ATP-treated neutrophils remained viable but still exhibited IL-1β release (**Fig 1G**). Additionally, a lower concentration of nigericin in LPS-primed neutrophils did not cause cell death, but led to IL-1β release (**Fig 1F, G)**. These data together advocate differential pathways for cell death and IL-1β release in neutrophils. In contrast, release of tumour necrosis factor (TNF)-α was primarily stimulated by the LPS, but not further increased with nigericin or ATP treatment in neutrophils **(Fig S1D)**. Further, priming with zymosan A (a TLR2 ligand) also triggered similar IL-1β release and cell death, in response to nigericin **(Fig S1E, F)**, confirming pyroptosis in nigericin-treated neutrophils under varying priming conditions. Microscopic analysis of intracellular IL-1β showed LPS priming was essential for the nigericin-induced IL-1β response in neutrophils (**Fig 1H**). Time-dependent analysis also confirmed an increase in IL-1β secretion with nigericin in the presence of LPS priming, but not without priming (**Fig 1I**). Furthermore, Calcien AM, a viability dye, disappeared in nigericin-treated cells independent of priming, but was retained in LPS+ATP treated neutrophils (**Fig 1J, K)**. Interestingly, Calcien AM disappeared before cellular swelling, which was followed by an increase in PI uptake in nigericin-treated neutrophils, confirming that different events occurred prior to plasma membrane integrity loss (**Fig 1L**). Together, our data identified distinct pyroptotic responses, including IL-1β release in LPS-primed ATP-treated live neutrophils, both cell death and IL-1β release in LPS-primed nigericin-treated neutrophils, and cell death without IL-1β and TNF-α release in nigericin-treated neutrophils (**Fig 1M**), providing a unique prospect to unravel the differential events and signaling involved in pyroptotic processes in neutrophils.

### NLRP3 and caspase-1/11 regulate IL-1β, while caspase-7/9 control pyroptotic cell death

Pyroptosis can be initiated by canonical or non-canonical inflammasome pathways via caspase-1 and caspase-11 respectively. These caspases cleave GSDMD, leading to the plasma membrane pore formation that precedes cell death (5). Interestingly, we observed a distinct morphological phenotype displayed by pyroptotic neutrophils, compared to NETotic and apoptotic deaths in neutrophils (**Fig 2A, B)**. Further, we recognize pyroptotic responses with the expression of NLRP3, cleaved caspase-1, and IL-1β release (**Fig 2C**). LPS-primed nigericin-treated neutrophils exhibited increased expression of NLRP3, caspase-1 activation, and mature IL-1β release (**Fig 2C-F**), while a lower level of increase was observed in LPS-primed ATP-treated neutrophils. Surprisingly, neutrophils treated with nigericin alone did not display NLRP3 and caspase-1 activation (**Fig 2C-F**). NETotic (Ionomycin and PMA treated) and apoptotic (Staurosporine treated) neutrophils did not exhibit activation of pyroptotic signaling components, including NLRP3, caspase-1, and IL-1β release (**Fig 2C-E**). Moreover, MCC950 (NLRP3 inhibitor) and VX765 (caspase-1 inhibitor) (25, 26), reduced IL-1β secretion in LPS-primed nigericin-treated neutrophils (**Fig 2G**), confirming the role of NLRP3 inflammasome and caspase-1 in IL-1β secretion. Conversely, LDH (140 KDa) release was alleviated with caspase-1 inhibitor, VX765, but not with MCC950 (**Fig 2H**) (27). These results further suggest the involvement of caspase-1 signaling in downstream pyroptotic cell death. Furthermore, caspase-11 activation with LPS and LPS+ATP treatments suggests the role of NLRP3/caspase-11 signaling in LPS+ATP mediated IL-1β secretion (**Fig 2I and Fig S2A)** (14, 21).

**Fig 2:**
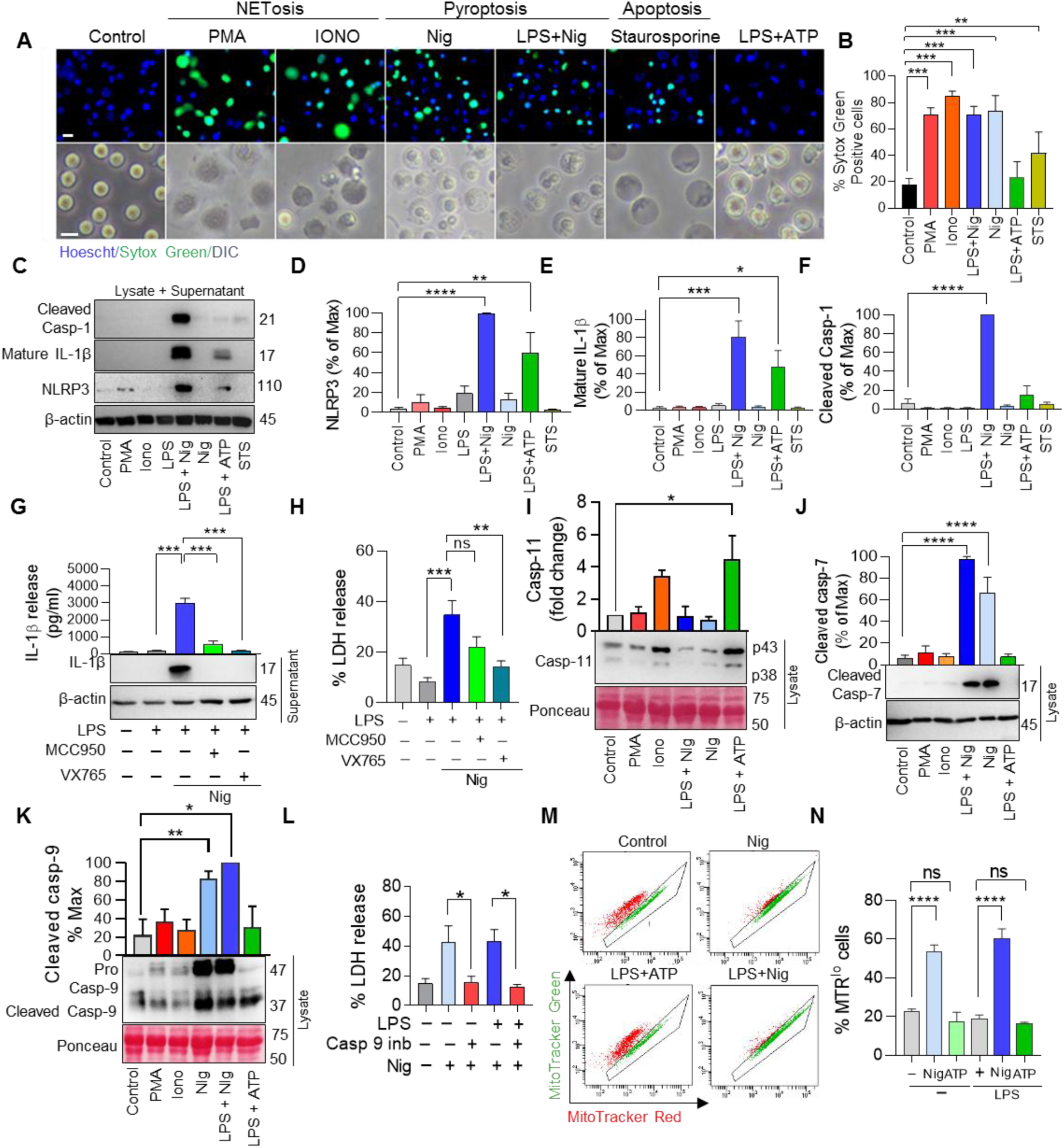
Analysis of signalling components, NLRP3/Caspase-1/ IL-1β in nigericin-induced neutrophil pyroptosis. **(A)** Representative image showing Hoechst (blue), Sytox Green (green) positive cells (Scale bar, 20 μm) and DIC (gray) (Scale bar, 10 μm) with different inducers for NETs (PMA, Ionomycin), Pyroptosis (nigericin, LPS + nigericin, LPS + ATP) and apoptosis (Staurosporine (STS)) in neutrophils. **(B)** Quantification showing Sytox green positive cells with different inducers (**P <0.01, ***P <0.001, n = 3 independent experiments, analyzed using One-way ANOVA). **(C)** Western blot image of NLRP3, cleaved caspase-1, and mature IL-1β release with different inducers. **(D)** Quantification of NLRP3 with different death inducers (**P <0.01, ****P <0.0001, n = 3 independent experiments, analyzed using One-way ANOVA). **(E)** Quantification of mature IL-1β release with different death inducers (*P <0.05, ***P <0.001, n = 3 independent experiments, analyzed using One-way ANOVA) **(F)** Quantification of cleaved caspase-1 with different inducers (****P <0.0001, n = 3 independent experiments, analyzed using One-way ANOVA). **(G)** IL-1β secretion in LPS + nigericin stimulated neutrophil in the presence or absence of MCC950 (NLPR3 inhibitor – 30μM), VX765 (caspase-1 inhibitor – 30μM) (***P <0.001, n = 4 independent experiments, analyzed using One-way ANOVA). **(H)** % LDH release in LPS + nigericin stimulated neutrophil in the presence or absence of MCC950, VX765 (**P <0.01, ***P <0.001, n = 4 independent experiments, analyzed using One-way ANOVA). **(I)** Representative western blot image and quantification showing caspase-11 activation with different inducers (*P <0.005, n = 3 independent experiments, analyzed using One-way ANOVA). **(J)** Western blot image and quantification of cleaved caspase-7 in neutrophils stimulated with different death inducers (****P <0.0001, n = 3 independent experiments, analyzed using One-way ANOVA). **(K)** Western blot image and quantification of pro-caspase-9 and cleaved caspase-9 expression in neutrophils stimulated with different death inducers (*P<0.05, **P <0.01, n = 3 independent experiments, analyzed using One-way ANOVA). **(L)** % LDH release in the presence of caspase-9 inhibitor (50 μM) with nigericin in non-primed and LPS-primed neutrophils (*P<0.05, n = 3 independent experiments, analyzed using One-way ANOVA). **(M)** Dot plots showing MitoTracker red and MitoTracker green staining in nigericin and ATP challenge in non-primed and LPS-primed neutrophils. **(N)** Quantification of the percentage of MitoTracker red low cells in non-primed and LPS-primed nigericin and ATP challenged neutrophils (****P <0.0001, n = 4 independent experiments, analyzed using One-way ANOVA).

The absence of caspase-1 and NLRP3 in cytokine uncoupled pyroptosis triggered by nigericin treatment prompted us to explore the underlying mechanism of cell death. Activation of caspase-7 in nigericin-treated and LPS-primed nigericin-treated neutrophils confirmed programmed cell death (**Fig 2J**). Additionally, increased pro-caspase and cleaved caspase-9 expression was observed in nigericin-treated neutrophils, independent of LPS priming (**Fig 2K**). Treatment with a caspase-9 inhibitor reduced LDH release in nigericin-treated neutrophils regardless of LPS priming, confirming the involvement of caspase-9 in nigericin-induced pyroptotic cell death (**Fig 2L**). In conjunction, plasma-membrane rupture requires pathways leading to the activation of caspase-9 and caspase-7 in murine neutrophils. These results demonstrate unique functions for caspase-1, caspase-11, caspase-9, and caspase-7 in neutrophil pyroptotic processes. We further investigated mitochondrial damage, which is linked to the aforementioned caspases and presents both a cause and consequence facet for inflammasome activation and pyroptosis (28, 29). Untreated and LPS-primed ATP-treated neutrophils displayed good mitochondrial health, as indicated by high mitochondrial potential analysed using mitochondrial membrane potential independent MitoTracker Green-FM and dependent MitoTracker Red CMXRos dyes (**Fig 2M, N)**. In contrast, nigericin consistently caused a loss of mitochondrial potential, independent of LPS priming (**Fig 2M, N)**. MitoSOX staining demonstrated significantly increased mitochondrial ROS in neutrophils upon nigericin treatment, indicating mitochondrial stress **(Fig S2B, C)**. Kinetic analysis of JC-1 dye showed a red tubular interconnected network presenting healthy mitochondria in the control group. Conversely, nigericin treatment led to a collapse of JC-1 red signal, representing loss of mitochondrial membrane potential, after 60 min **(Fig S2D, E)**. Subsequently, JC-1 green signal increased concurrently, demonstrating the loss of active, healthy mitochondria **(Fig S2D, E)**. Intriguingly, the JC-1 green signal collapsed, followed by cellular swelling, increased membrane permeability, and eventual plasma membrane rupture **(Fig S2D, E)**. These data suggested mitochondrial damage is a prerequisite for neutrophil viability collapse during pyroptosis, but mitochondrial signaling modulators, including rotenone (a mitochondrial complex I inhibitor), antimycin (a mitochondrial complex III inhibitor), and FCCP (a mitochondrial uncoupler) did not significantly alter LPS-primed nigericin-induced IL-1β secretion **(Fig S2F)**. Oligomycin (an inhibitor of mitochondrial complex V or ATP synthase) provided some protection, possibly driven by purinergic signaling in inflammasome activation (20). Interestingly, LDH release was not mitigated by these mitochondrial modulators **(Fig S2G)** (30). These findings are consistent with previous observations describing the minimal role of mitochondria in neutrophil energetics and functions.

### Pyroptotic cell death exhibits nuclear rounding, cell swelling, GSDMD-dependent pore formation and cell rupture in neutrophils

We next inspected cellular and morphological features associated with distinct pyroptotic fates observed in neutrophils. Nigericin treatment caused cellular swelling/ ballooning, loss of plasma membrane integrity, independently of LPS priming in neutrophils (**Fig 3A, B, S3A)**. While LPS+ATP-treated cells exhibited morphology similar to the control group (**Fig 3A, B)**. Time-kinetic analysis in nigericin-treated neutrophils exhibited nuclear rounding, starting at early time points of around 30-90 min, followed by cell swelling/ ballooning at 90-120 min, and loss of plasma membrane integrity (**Fig 3C, D)**. These observations were further confirmed using flow cytometric analysis showing an initial increase in cell size (FSC), followed by nuclear reorganization (Hoechst intensity), increase in granularity (SSC), and later plasma membrane integrity loss (Sytox green positivity) (**Fig 3E**). An increase in intensity of Hoechst-33342 has been reported during nuclear condensation (31). We observed distinct stages of pyroptotic cell death, including cells displaying increased nuclear rounding, swelling/ballooning without Sytox Green uptake **(Fig S3A)**. Cells with swelling, a round nucleus, and Sytox Green positivity represented a later stage **(Fig S3A)**. Additionally, a fraction of small-sized cells with Sytox Green positive round nucleus, suggested loss of cellular content with cell rupture **(Fig S3A)**. Indeed, time lapse analyses further confirmed cell rupture by a collapse in size **(Fig S3B)**, and forward scatter (**Fig 3E**) following cell swelling.

**Fig 3:**
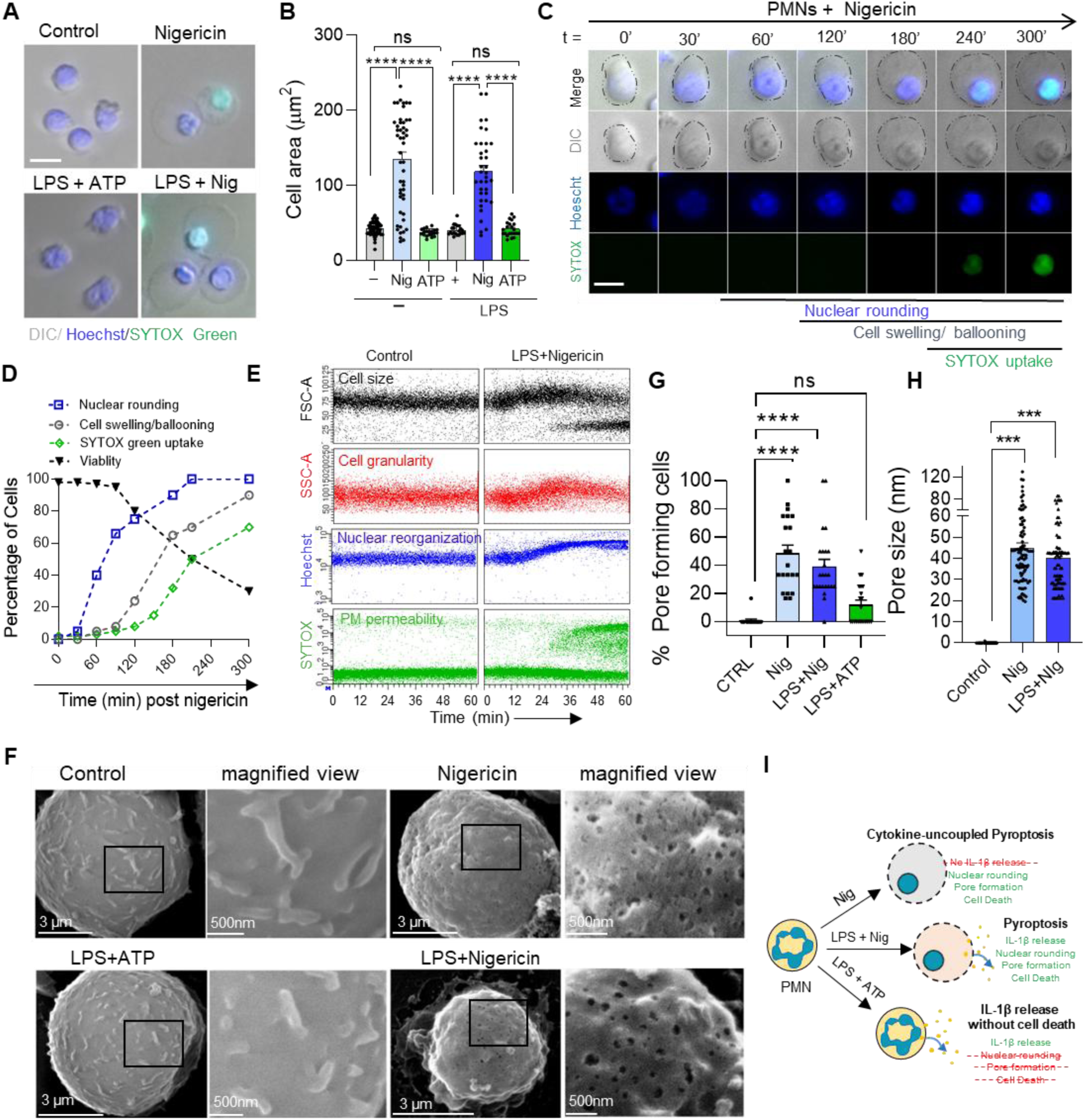
Pyroptosis proceeds via nuclear rounding, cell swelling, followed by loss of plasma membrane permeability in neutrophils. **(A)** Representative image of DIC, Hoechst, and Sytox green staining in neutrophils after nigericin and ATP treatment with and without LPS priming in neutrophils (Scale bar, 10 μm). **(B)** Quantification of cell area (μm^2^) after nigericin and ATP treatment with and without LPS priming in neutrophils (****P <0.0001, Data analysed from >500 cells in n=3 independent experiments, analyzed using One-way ANOVA). **(C)** Representative image showing time-dependent analysis of the sequence of pyroptotic events after nigericin challenge in murine neutrophils using DIC (gray), Hoechst (blue) and Sytox Green (green) (Scale bar, 5 μm). **(D)** Quantification showing time-dependent analysis for the sequence of pyroptotic events after nigericin challenge in murine neutrophils. **(E)** Time kinetics for FSC-A (black), SSC-A (red), Hoechst (blue), and Sytox Green (green) mean fluorescence intensity in control and LPS + nigericin treated neutrophils analysed using flow cytometry. **(F)** Scanning electron micrographs of nigericin, LPS + nigericin and LPS + ATP-treated cells. The right-side panel indicates a higher magnification of boxed area, marked in cells on the left-side with different treatments. **(G)** Quantifications showing % percent pore-forming cells in nigericin, LPS + nigericin, and LPS + ATP-treated cells (****P <0.0001, ns-non-significant, n=2 independent experiments with a minimum of 40 cells per experiment, analyzed using One-way ANOVA). **(H)** Pore size (nm) in nigericin, LPS + nigericin, and LPS + ATP-treated cells (***P <0.001, ns-non-significant, n=2 independent experiments with a minimum of 40 cells per experiment, analyzed using One-way ANOVA). **(I)** Schematic diagram showing distinct morphological phenotypes observed with LPS+ Nig, Nig, and LPS+ATP treatments.

The plasma membrane pore formation is a hallmark of pyroptosis (32). High-resolution scanning electron microscopy analysis of nigericin-treated neutrophils exhibited numerous pores (ranging from 20-100 nm in diameter) on the plasma membrane, independent of LPS priming (**Fig 3F-H**). Furthermore, inhibition of GSDMD through necrosulfonamide (NSA) prevented LPS+Nig induced IL-1β and LDH release in neutrophils, confirming involvement of GSDMD-dependent pores in regulating these outcomes **(Fig S3C, D)** (10, 11, 19). As expected, control neutrophils showed numerous microvilli on their surface (**Fig 3F**). Further, negligible pores were observed in LPS-primed ATP-treated neutrophils suggesting other distinct mechanisms regulating the IL-1β release (**Fig 3F, G)**. This aligned with the previous study, indicating the absence of plasma membrane pore formation in neutrophils (6). Altogether, neutrophils exhibit distinct events including nuclear rounding, cell swelling, and pore formation leading to pyroptotic cell death with nigericin treatment independent of LPS priming (**Fig 3I**).

### Enhanced DNA damage instigates neutrophil pyroptotic death but reduced IL-1β secretion

Our data so far indicate that LPS priming is essential for inflammasome activation and IL-1β processing in neutrophils, while it is dispensable for nigericin-dependent cell swelling and pyroptotic death. Interestingly LPS-priming induces extracellular signal-regulated kinase 1/2 (ERK1/2) and Protein kinase B (PKB)/Akt activation, that can regulate inflammasome activation and survival in neutrophils (33–35). We observed heightened and sustained Akt activation in LPS-primed nigericin and ATP-treated neutrophils (**Fig 4A, B)**. ERK1/2 activation was most prominent in LPS-primed ATP-treated neutrophils, whereas LPS-primed nigericin-treated neutrophils exhibited intermediate activation levels (**Fig 4A, C)**. Conversely, treatment with nigericin alone displayed diminished Akt and ERK1/2 activation (**Fig 4A-C**). This suggested that LPS-priming is crucial for Akt and ERK1/2 activation, while compromised cell survival signaling leads to death in nigericin-treated neutrophils. To further elucidate their involvement in the regulation of pyroptotic responses, we utilized pharmacological inhibitors of PI3K/AKT/MAPK signalling, including wortmannin, MK2206, and SB203580 (35). Inhibition of mitogen-activated protein kinase (MAP kinases) did not affect IL-1β secretion or LDH release **(Fig S4A, C)**. However, it led to a reduction in TNF-α levels **(Fig S4B)**, consistent with previous studies showing the involvement of Akt and p38 kinases, but not PI3K, in regulating TNF-α secretion (36). This indicates independence of these pathways for pyroptosis and IL-1β secretion, but engagement in inflammatory cytokine secretion.

**Fig 4:**
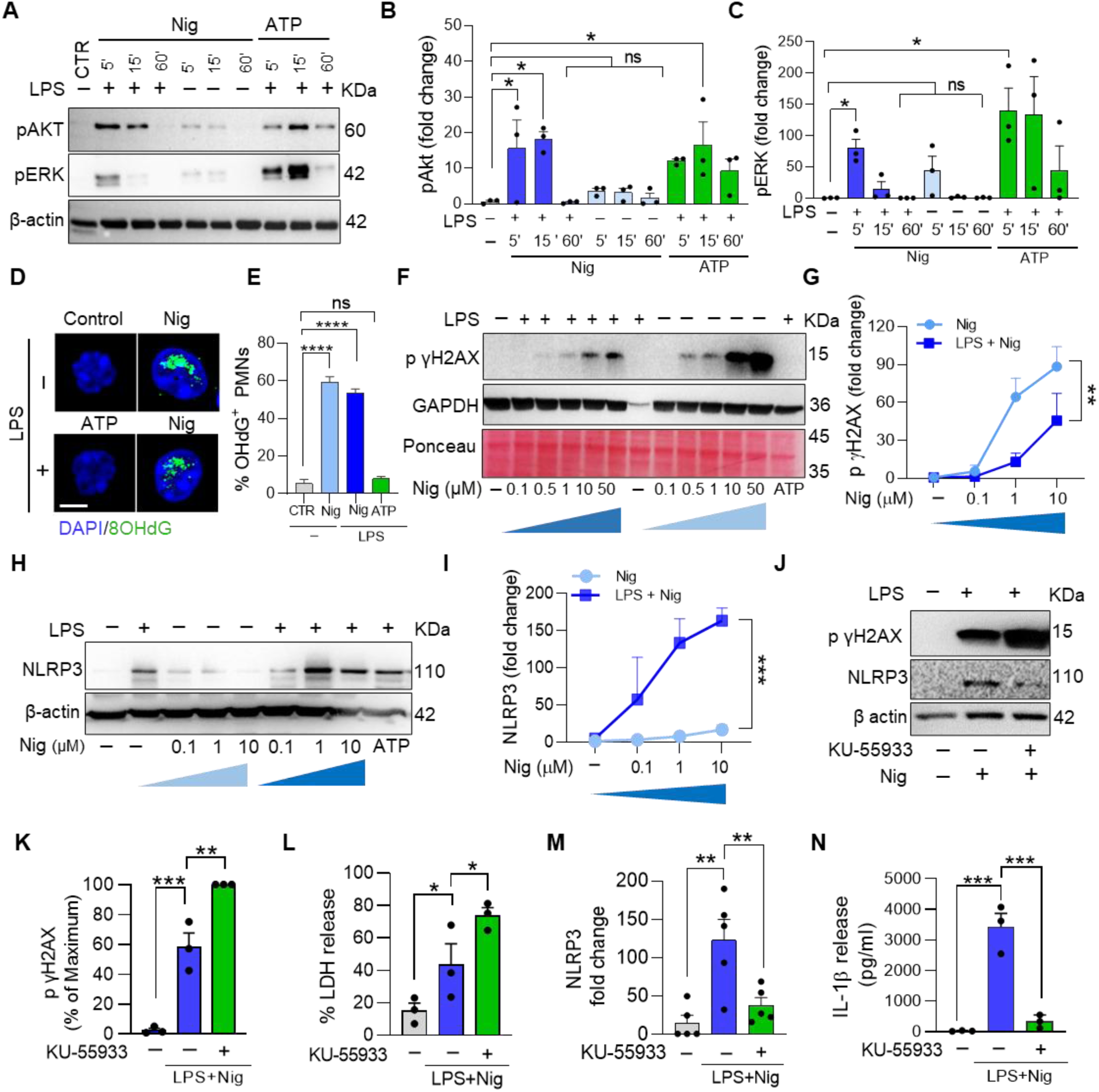
Reduced Akt and ERK signalling and enhanced DNA damage derived nigericin-induced pyroptotic death. **(A)** Western blot image showing pAkt and pERK expression in neutrophils after 5’, 15’ and 60’ of nigericin and ATP addition in non-primed or LPS-primed neutrophils. **(B)** Relative fold change of pAkt expression in neutrophils after 5’, 15’, and 60’ of nigericin and ATP addition in non-primed or LPS-primed neutrophils (*P<0.05, n = 3 independent experiments, analyzed using One-way ANOVA). **(C)** Relative fold change of pERK expression after 5’, 15’, and 60’ of nigericin and ATP addition in non-primed or LPS-primed neutrophils (*P<0.05, n = 3 independent experiments, analyzed using One-way ANOVA). **(D)** Representative immunofluorescence image stained for 8-OHdG (green) and nucleus (blue) in nigericin or ATP-treated neutrophils (Scale bar, 5 μm). **(E)** Quantification of 8-OHdG positive cells after nigericin or ATP challenge in LPS primed neutrophils (****P <0.0001, n =2 independent experiments, analyzed using One-way ANOVA). **(F)** Western blot image showing dose-dependent effect of nigericin on p γH2AX expression with and without LPS priming in neutrophils. **(G)** Quantification of dose-dependent effect of nigericin on p γH2AX in the presence and absence of LPS priming (**P <0.01, n =3 independent experiments). **(H)** Western blot image showing NLRP3 expression in nigericin-treated neutrophils in the presence and absence of LPS priming. **(I)** Quantification of NLRP3 expression in nigericin-treated neutrophils in the presence and absence of LPS priming (***P <0.001, n = 3 independent experiments, analyzed using unpaired t-test). **(J)** Western blot image of p γH2AX and NLRP3 expression in LPS + nigericin-treated neutrophils in the presence and absence of KU-55933 (ATM kinase inhibitor, 10 μM). **(K)** Quantification of p-γH2AX in LPS + nigericin-treated neutrophils in the presence and absence of KU-55933 (**P <0.01, ***P <0.001, n = 3 independent experiments, analyzed using One-way ANOVA). **(L)** % LDH release in LPS + nigericin treated neutrophils in the presence and absence of KU-55933 (*P<0.05, n = 3 independent experiments, analyzed using One-way ANOVA). **(M)** Quantification of fold change in NLRP3 expression relative to control in LPS + nigericin treated neutrophils in the presence and absence of KU-55933 (**P <0.01, n = 5 independent experiments, analyzed using One-way ANOVA). **(N)** Quantification of IL-1β release in supernatant of LPS + nigericin treated neutrophils in the presence and absence of KU-55933 (***P <0.001, n = 5 independent experiments, analyzed using One-way ANOVA).

Based on the loss of nuclear lobulation following nigericin treatment preceding pyroptotic death, we hypothesized a link between DNA damage and nuclear rounding. Nigericin treatment significantly increased oxidized nucleotide 8-Hydroxy-2′-deoxyguanosine (8-OHdG) in neutrophils, indicating oxidative DNA damage (**Fig 4D, E)**. DNA damage was validated using another marker, histone γH2AX, that marked double-stranded DNA breaks. Nigericin stimulation resulted in a dose-dependent increase in phosphorylated γH2AX, while interestingly pre-priming with LPS protected cells from nigericin-induced γH2AX activation (**Fig 4F, G)**. Furthermore, LPS primed nigericin-treated neutrophils exhibited enhanced NLRP3 expression in a concentration and time-dependent manner (**Fig 4H, I and Fig S5A-C**). While no NLRP3 expression was observed with nigericin treatment (**Fig 4H, I and Fig S5A-C**), consistent with a previous report (37). Hence, LPS-priming acts responsible for the differential DNA damage and inflammasome signaling cascades. It is pertinent to mention that control and LPS-primed ATP-treated cells did not show any γH2AX activation **(Fig S5D)**.

Previous studies have shown activation of DNA damage response pathways, specifically Ataxia-telangiectasia mutated (ATM) regulates the secretion of various pro-inflammatory cytokines, including IL-1β release (38). We reasoned the role of DNA damage in diminished cytokine release response in nigericin-treated neutrophils and employed KU-55933, an inhibitor of the DNA damage repair pathway, ATM kinase, to LPS-primed nigericin-treated neutrophils. Inhibition of the DNA damage repair pathway led to an enhanced p-γH2AX expression (**Fig 4J, K)**. Moreover, this increased DNA damage was associated with elevated LDH release (**Fig 4L**), but surprisingly NLRP3 inflammasome expression and IL-1β secretion were reduced (**Fig 4M, N)**. Altogether, increased DNA damage signaling inhibits NLRP3 expression and IL-1β secretion, while enhanced pyroptotic cell death in neutrophils.

### Perinuclear F-actin polymerization and cytoskeletal reorganization facilitates nuclear rounding during pyroptosis

Increased DNA damage signalling and nuclear rounding appear to be major events in the regulation of IL-1β and cell death outcomes in neutrophils. Therefore, we further focused on the nuclear rounding phenomenon during pyroptosis. Untreated neutrophils exhibited a multi-lobulated and segmented nucleus (**Fig 5A**). In contrast, nigericin treatment significantly increased nuclear rounding independent of LPS priming in neutrophils (**Fig 5A, B)**. Multi-lobulated nuclear structures in neutrophils are regulated by the LINC (linker of nucleoskeleton and cytoskeleton) complex of nuclear membrane proteins, including lamins (Lamins A/C and B), Lamin B receptor (LBR), and cytoskeletal proteins (39). Moreover, cytoskeletal elements tether to the nuclear membrane, transducing external forces and signals to the nucleus (39). To understand the putative cytoskeletal reorganization, we evaluated F-actin organization in pyroptotic neutrophils exhibiting nuclear rounding. Control neutrophils with segmented nuclei displayed surface/cortical F-actin localization, but amazingly, a perinuclear actin ring structure was observed in nigericin-treated neutrophils displaying a round nucleus, which was also visualized with side view analysis of Z-stack images and line of interest analysis (**Fig 5C**). Moreover, control neutrophils displayed visible microvilli with cortical F-actin in the 3D reconstruction of cells (**Fig 5D**). Intriguingly, nigericin treatment substantially reduced F-actin staining and microvilli at the cell surface, while profoundly increased F-actin in the perinuclear region (**Fig 5D).** These data together suggested the formation of a F-actin cage or ring-like structure in providing prospective compressive force contributing to nuclear rounding.

**Fig 5:**
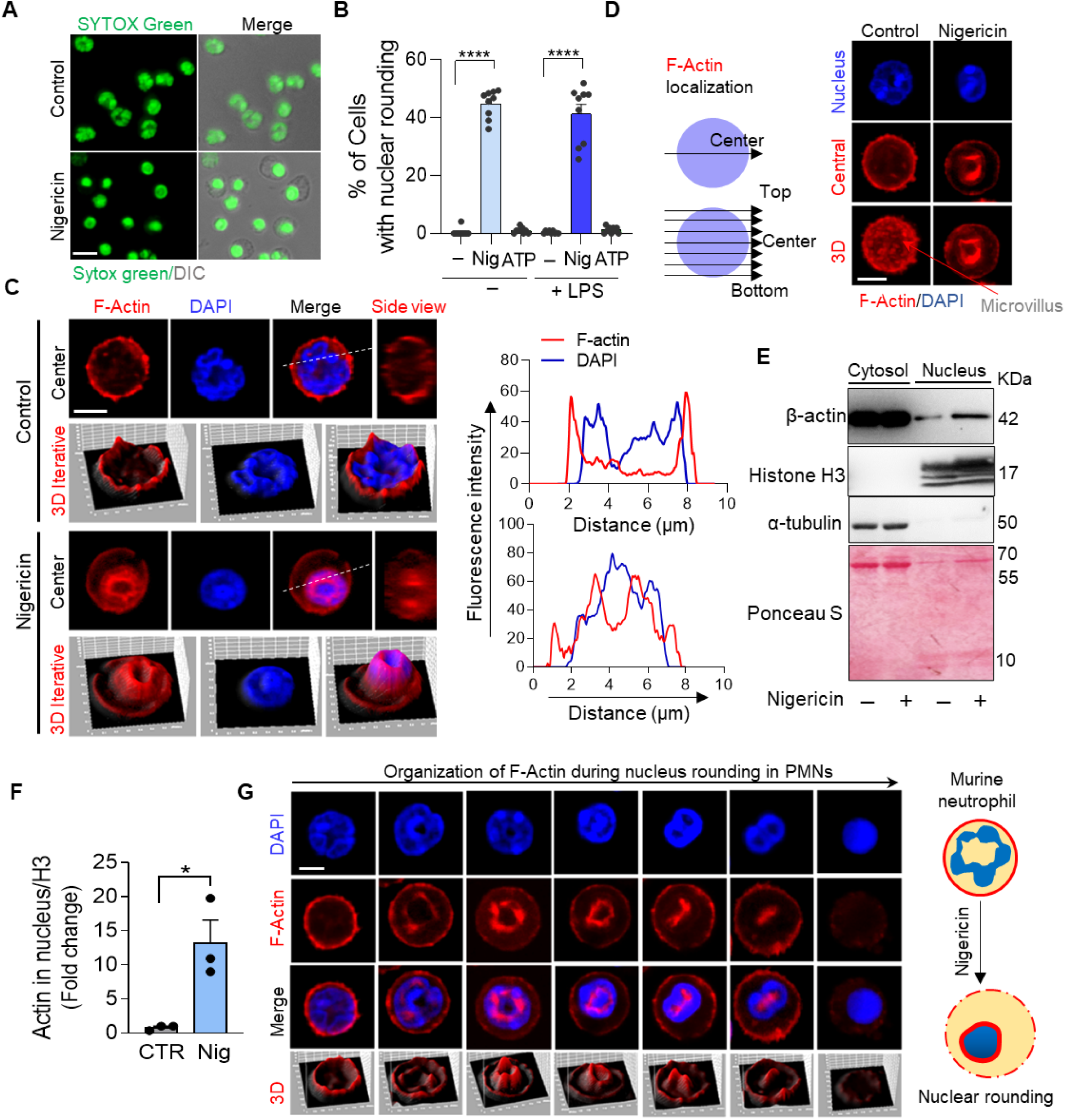
Perinuclear structural organization and F-actin polymerization facilitate nuclear rounding during pyroptosis in neutrophils. **(A)** Representative image stained for SYTOX green showing nuclear changes in LPS-primed nigericin-treated PMNs (Scale bar, 10 μm). **(B)** Quantification of percent cells showing nuclear rounding in nigericin and ATP-treated PMNs with and without LPS-priming (****P <0.0001, n = 8 independent experiments, analyzed using One-way ANOVA). **(C)** Representative confocal image and line intensity profile showing side view, central region and 3D construct indicating F-actin perinuclear localization in nigericin-stimulated neutrophils stained with rhodamine phalloidin (red) and DAPI (blue) (n = 2 independent experiments, with a minimum of 200 cells per experiment analyzed) (Scale bar, 5 μm). **(D)** Control showing microvilli with cortical F-actin content upon 3D construction, while actin ring in the perinuclear region of nigericin-treated neutrophils (Scale bar, 5 μm) (n = 2 independent experiments). **(E)** Representative western blot image showing actin levels in cytosolic vs nuclear fraction with and without nigericin treatment in neutrophils. **(F)** Quantification showing fold change in actin levels in nuclear fraction of untreated and nigericin-treated neutrophils (*P<0.05, n = 3 independent experiments, analysed using unpaired t test). **(G)** Representative image stained with rhodamine phalloidin (red) and DAPI (blue) showing F-actin localization in different stages of nuclear rounding (n = 2 independent experiments, with a minimum of 150 cells per experiment analyzed) (Scale bar, 5 μm).

The microscopic observation of F-actin association with nucleus was validated using cytosolic and nuclear fractionation. The nuclear fraction of nigericin-treated neutrophils confirmed an increase in actin content compared to the control neutrophils (**Fig 5E, F)**. Histone and tubulin confirmed the nuclear and cytosolic fractionation respectively (**Fig 5E**). We further evaluated the crosstalk between F-actin and nucleus in neutrophils at different stages of nuclear rounding (**Fig 5G**). The data indicated the early development of the F-actin ring in neutrophils displaying a relaxed and lobulated nucleus, which became more prominent as the nucleus displayed rounding. Subsequently, these neutrophils displayed a linear F-actin structure likely pulling nuclear lobules. Finally, cells exhibiting complete rounding of the nucleus presented diminished f-actin staining (**Fig 5G**), likely degradation by caspase-dependent mechanisms and death at later time point.

### Rho kinase signalling regulates nigericin-induced nuclear rounding and pyroptotic cell death

Based on the accumulation of F-actin in the nucleus, we further investigated the presence of actin-associated motor protein, myosin light chain (MLC), as a putative contractile force during nuclear rounding in nigericin-treated neutrophils. 3-D analysis revealed colocalization of the F-actin ring with an active form of myosin, i.e. phosphorylated MLC (pMLC) protein in the perinuclear region of nigericin-treated neutrophils (**Fig 6A, B)**. These findings suggested the involvement of cytoskeletal reorganization in the regulation of nuclear rounding during pyroptosis.

**Fig 6:**
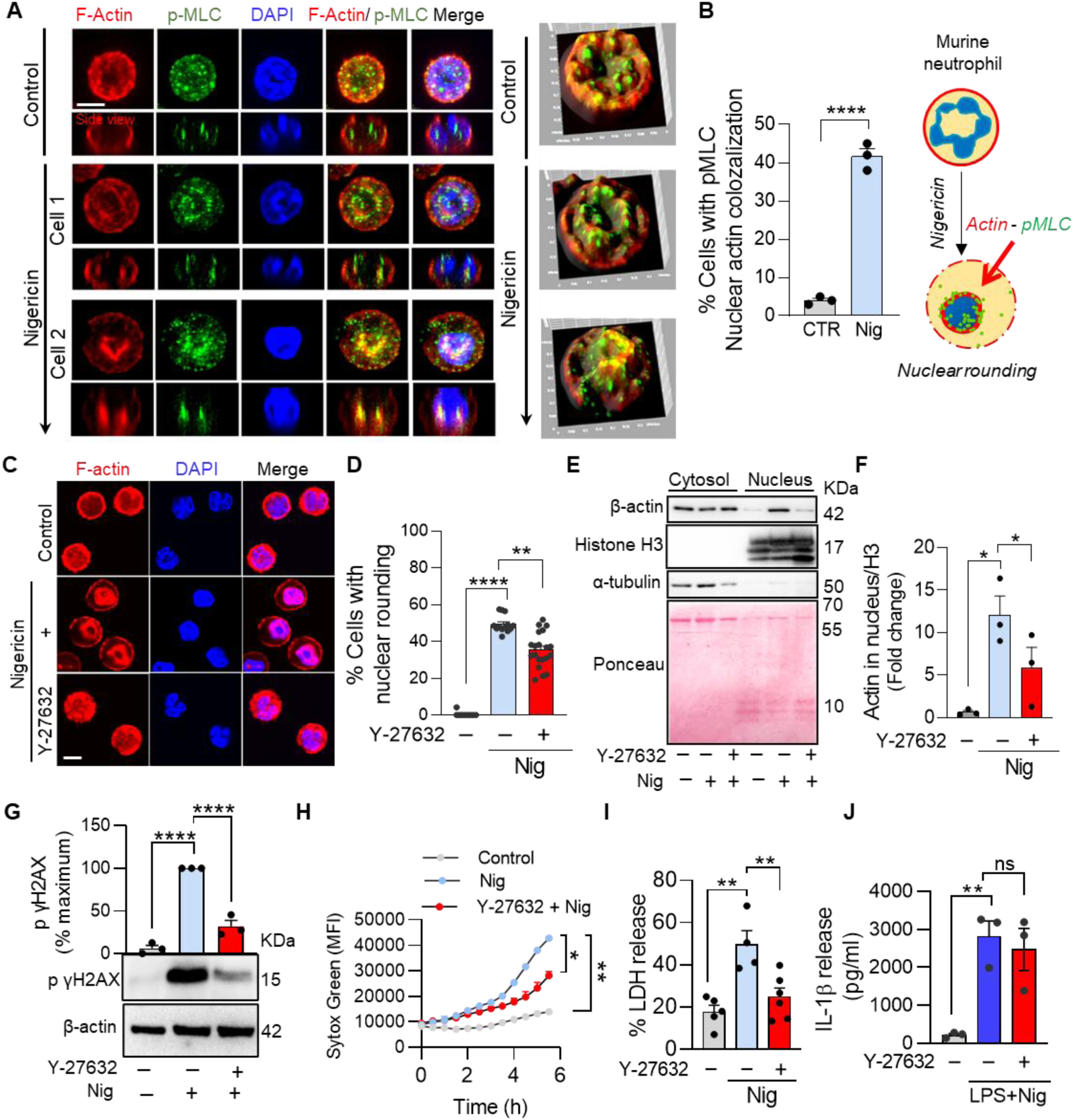
Actomyosin signaling is critical for nuclear rounding and cell death for neutrophil pyroptosis. **(A)** Representative confocal image of LPS + nigericin-treated neutrophils stained for pMLC (green), F-actin (red), and nucleus (blue) (Scale bar, 5 μm). **(B)** Quantification showing percentage of cells with pMLC – nuclear actin colocalization in nigericin-treated PMNs (****P <0.0001, n = 2 independent experiments, with a minimum of 150 cells per experiment analyzed, analysed using paired t-test). **(C)** Representative image showing F-actin (red) and nuclear rounding (blue) with rho-kinase inhibitor Y-27632 (30μM) in nigericin-treated neutrophils (n = 2-4 independent experiments) (Scale bar, 5 μm). **(D)** Relative quantification of % cells showing nuclear rounding in the presence of Y-27632 (30μM) in nigericin-treated neutrophils (**P <0.01, ****P <0.0001, n = 4 independent experiments, with a minimum of 100 cells per experiment analyzed, analysed using One-way ANOVA). **(E)** Western blot image showing actin expression in the nuclear fraction of nigericin-treated neutrophils and its rescue with Y-27632 (30μM). **(F)** Quantification showing actin expression in the nuclear fraction of nigericin-treated neutrophils and its rescue with Y-27632 (30μM) (*P<0.05, n = 3 independent experiments, analysed using paired t-test). **(G)** Western blot image and quantification of p yH2AX with nigericin in the presence or absence of Y-27632 (30μM) (****P <0.0001, n = 3 independent experiments, analysed using One-way ANOVA). **(H)** Time-dependent Sytox Green uptake with LPS+nigericin in the presence or absence of Y-27632 (30μM) (*P <0.05, **P <0.01, n = 3 independent experiments, analysed using One-way ANOVA). **(I)** % LDH release with LPS+nigericin in the presence or absence of Y-27632 (30μM) (**P <0.01, n = 3 independent experiments, analysed using One-way ANOVA). **(J)** IL-1β release (pg/ml) with LPS+nigericin in the presence or absence of Y-27632 (30μM) (**P <0.01, n = 3 independent experiments, analysed using paired t-test).

During pyroptosis, neutrophils treated with nigericin exhibit distinct nucleus shape transitioning from ring/lobulated to round **(Fig S6A, B)**. To investigate the involvement of the cytoskeleton and other regulatory pathways in nuclear rounding, we employed various pharmacological inhibitors and quantified cells displaying distinct nuclear phenotypes **(Fig S6B)**. Remarkably, targeting cytoskeletal reorganization and actomyosin contractility with the Rho-associated protein kinase (ROCK) inhibitor, Y-27632, significantly abrogated nigericin-induced nuclear rounding and protected the segmented nucleus, confirming the role of cytoskeletal reorganization (**Fig 6C, D and Fig S6B)**. Consistently, microscopy data showed Y-27632 treatment prevented nigericin-induced perinuclear actin ring formation in the nucleus (**Fig 6C**), which was further validated biochemically with actin levels in the nuclear fraction (**Fig 6E, F)**. Additionally, inhibition of caspases with pan-caspase inhibitor, Q-VD-OPh, and caspase-1 inhibitor, VX765 significantly abrogated nigericin-induced nuclear rounding in neutrophils, indicating the role of caspases in the upstream signaling for nuclear rounding **(Fig S6B)**. This aligns with the standing evidence showing caspases participating in the DNA degradation processes (40). Conversely, NLRP3 inflammasome inhibitor, MCC950, did not affect nigericin-induced nuclear rounding **(Fig S6B)**, suggesting NLRP3-independent reorganization of nuclear structure. Moreover, inhibition of DNA repair via ATM Kinase inhibitor, KU-55933 enhanced nigericin-induced nuclear rounding and DNA damage **(Fig S6B and Fig 4J, K)**. These data correlated with protected γH2AX phosphorylation, and inhibited DNA damage observed in Y-27632, Q-VD-OPh, and VX765 treated neutrophils (**Fig 6G and S6C)**. Y-27632 mediated protection was well supported by the decrease in Sytox Green uptake and LDH release in LPS-primed nigericin-stimulated neutrophils (**Fig 6H, I)**, while no effect was observed on IL-1β release (**Fig 6J**). This suggested a specific and significant role of actomyosin reorganization and DNA damage in the neutrophil pyroptotic death process, rather than in IL-1β release.

### Bacteria-induced pyroptosis displays nuclear rounding, cell swelling, and IL-1β phenotype

To further validate the relevance of the above observations, we utilized a bacteria-induced pyroptosis model in neutrophils (8). Morphological analyses of neutrophils challenged with *Escherichia coli (E. coli)* revealed increased cell swelling, a pyroptotic death feature (**Fig 7A**). The LDH release and IL-1β secretion upon *E.coli* exposure established bacteria-induced pyroptotic cell death in neutrophils (**Fig 7B, C)**. Immunofluorescence imaging confirmed nuclear rounding and perinuclear localization of F-actin in *E. coli* challenged neutrophils (**Fig 7D**). This phenomenon was corroborated by increased actin expression in the nuclear fraction of *E.coli* treated neutrophils (**Fig 7E**). These data recapitulate the nuclear rounding phenotype in bacteria-induced pyroptosis, suggesting the uniformity of this phenomenon in neutrophil pyroptosis under distinct settings.

**Fig 7:**
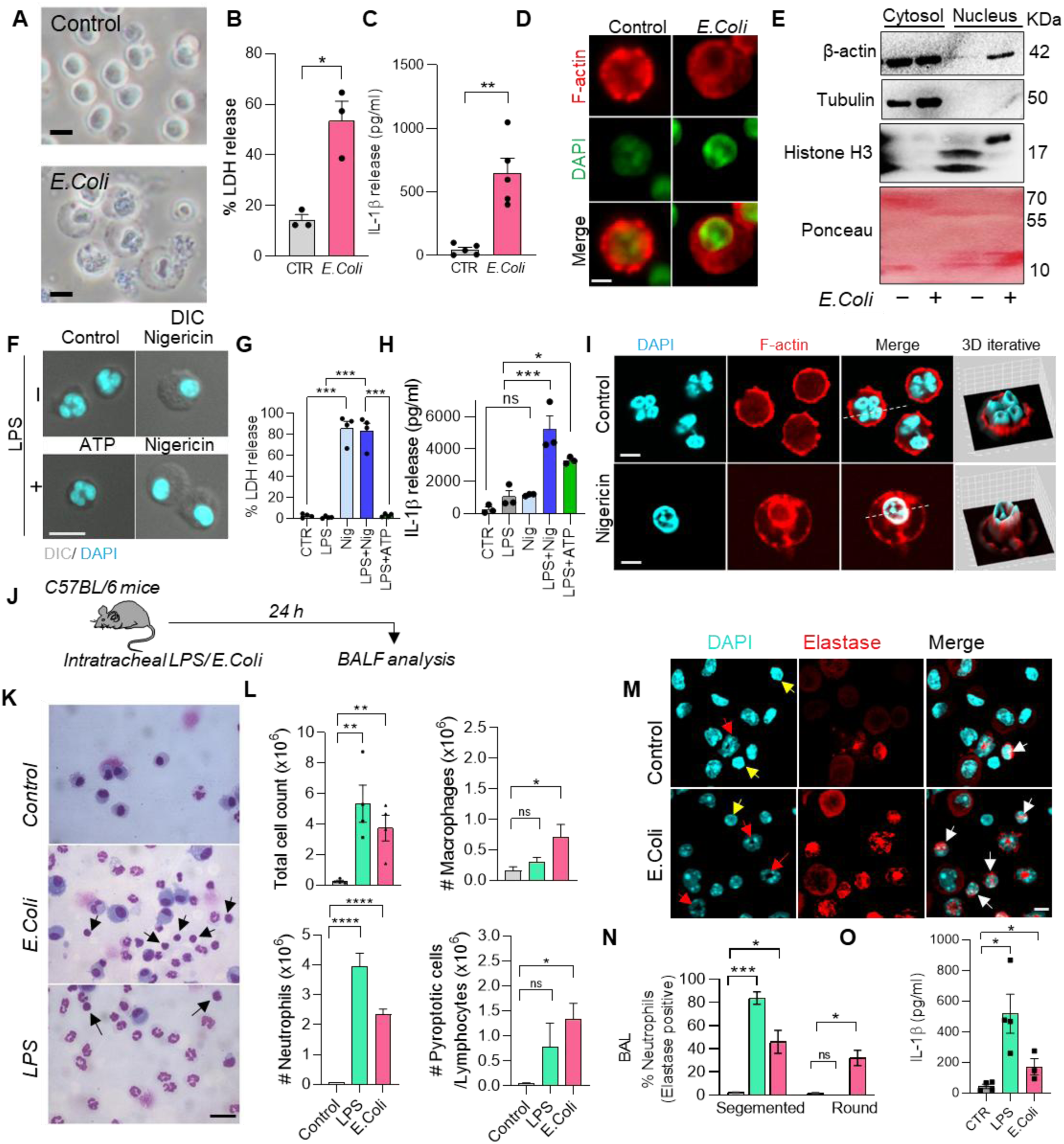
Cytoskeletal and nuclear reorganization during pyroptosis in different conditions, including human neutrophils. **(A)** Representative image showing morphological changes showing nuclear rounding, cell swelling in murine neutrophils upon *E.coli* infection using DIC (n = 2 independent experiments). **(B)** % LDH release in neutrophils infected with *E.coli* at MOI of 10 (*P<0.05, n = 3 independent experiments, analyzed using paired t test). **(C)** IL-1β release in neutrophils infected with *E.coli* at MOI of 10 (**P<0.01, n = 5 independent experiments, analyzed using paired t test). **(D)** Immunofluorescence image showing nuclear rounding (green) and F-actin ring (red) in the perinuclear region of *E. coli-treated* PMNs (n = 3 independent experiments) (Scale bar, 5 μm). **(E)** Representative western blot showing actin expression in the nuclear fraction of control and *E. coli-treated* murine neutrophils. **(F)** Representative DIC image (gray) and Hoechst (cyan) staining showing cell swelling and nuclear rounding in human neutrophils in response to nigericin treatment with and without LPS priming. (n = 3 independent experiments, with a minimum of 400 cells per experiment analyzed) (Scale bar, 10 μm). **(G)** % LDH release with nigericin and ATP treatment with and without LPS priming (***P <0.001, n = 4 independent experiments, analysed using One-way ANOVA). **(H)** IL-1β release with nigericin and ATP treatment with and without LPS priming (*P<0.05, ***P <0.001, n = 3 independent experiments, analysed using One-way ANOVA). **(I)** Representative confocal image and 3D iterative analysis of Hoechst-positive nucleus (cyan) and Rhodamine-phalloidin (red) displaying F-actin in control and nigericin-stimulated neutrophils (Scale bar, 5 μm). **(J)** Schematic diagram showing the strategy of LPS and *E.Coli* acute lung injury model development in mice. **(K)** Representative Giemsa image of BAL cells in *E.Coli* and LPS-challenged mice. Black arrows indicate cells with a round nucleus (Scale bar, 20 μm). **(L)** Quantifications of total cell, neutrophil, macrophage, and Pyroptotic/lymphocyte count in BAL of *E.Coli* and LPS challenged mice (**P<0.01, ****P<0.0001, Data represent n = 4 mice). **(M)** Representative immunofluorescence image showing elastase-positive (red) neutrophils with segmented nucleus and nuclear rounding (cyan) in BAL cells of *E.Coli* and LPS challenged mice. Red arrows indicate elastase + cells with a segmented nucleus, yellow arrow marks the cells with a round nucleus without elastase, and white arrow marks the elastase + cells with a round nucleus (Scale bar, 5 μm). **(N)** Quantification of elastase + cells with a segmented nucleus or a round nucleus (***P<0.001, *P <0.05, Data represent n = 4 mice). **(O)** IL-1β levels (pg/ml) in the BALF of *E. Coli* and LPS challenged mice (*P <0.05, Data represent n = 4 mice, analysed using One-way ANOVA).

### Human neutrophils exhibit nuclear rounding and other pyroptotic characteristics

Remarkably, consistent with murine studies, we observed that human neutrophils exhibited pronounced nuclear rounding and cell swelling when exposed to nigericin, irrespective of LPS priming (**Fig 7F, S7A)**. While ATP did not elicit these phenotypes (**Fig 7F, S7A)**. Both Sytox permeability and LDH release studies confirmed cell rupture with nigericin independent of LPS priming, but not with ATP **(Fig S7B, 7 G)**. Intriguingly, LPS + ATP was sufficient to trigger human neutrophils to secrete IL-1β without pyroptotic cell death (**Fig 7H**). Moreover, LPS-primed nigericin-stimulated human neutrophils induced significant IL-1β secretion (**Fig 7H**). Interestingly similar to mice neutrophils, IL-1β secretion was not induced in human neutrophils in response to nigericin alone, confirming a prerequisite role of LPS priming for IL-1β secretion (**Fig 7H**). Furthermore, time-dependent analyses revealed a transition of highly lobulated neutrophils to round nuclei, without chromatin decondensation **(Fig S7C, D)**. Consistently, we observed F-actin on the cell surface of control neutrophils, which moved to the perinuclear region, forming a cage-like structure in response to nigericin, as observed in Z-stack and line of interest analyses (**Fig 7I**). Together, these data recapitulate distinct pyroptotic responses, including IL-1β release in LPS-primed ATP-treated neutrophils, both cell death with IL-1β release in LPS-primed nigericin-treated neutrophils, and cell death without IL-1β release in nigericin-treated human neutrophils, with likely similar mechanisms of DNA damage, cytoskeletal reorganization, caspase activation, and nuclear rounding observed in murine neutrophils.

### Acute lung injury model displays enhanced neutrophil infiltration, pyroptosis and nuclear rounding

To further look for in vivo relevance, we utilized acute lung injury models. Mice were challenged intratracheally with PBS, LPS or heat-inactivated bacteria (**Fig 7J**). Lungs and bronchoalveolar lavage fluid (BALF) were analyzed at 24 h to investigate both infiltrating cells and neutrophils exhibiting pyroptotic changes. The data revealed significant influx of leukocytes, majorly neutrophils (**Fig 7L**). Giemsa staining identified higher frequency of segmented neutrophils in LPS challenged mice, while heat-inactivated bacteria challenged mice displayed ∼50% of segmented neutrophils (**Fig 7K**). Of note, we observed 30-40% cells with roundish nucleus, to be either lymphocytes or neutrophils displaying round nucleus (**Fig 7K**). Immunofluorescence staining of elastase, a primary granule protein of neutrophils, further differentiates these cells and confirms most of these cells as elastase+ neutrophils with roundish nucleus (**Fig 7M, N)**. While only a minor fraction of cells were likely elastase negative lymphocytes. The analysis of BALF displayed enhanced IL-1β release in LPS and heat-inactivated bacteria challenged mice. These data together establish the relevance of neutrophil pyroptotic responses in inflammatory conditions.

## DISCUSSION

This study describes neutrophil pyroptotic responses under different conditions and dissects distinct pyroptosis-associated events, including context-dependent differential IL-1β release, nuclear rounding, cell swelling/ ballooning, and cell rupture in murine and human neutrophils. It is important to note that pyroptosis, which virtually means “fiery cell death,” is often associated with IL-1β release, cell swelling, pore formation, and plasma membrane rupture. Though recent research described independence of pyroptotic events under specific conditions, the term ‘’pyroptosis’’ continues to be referenced casually. Here, we described distinct outcomes observed, including 1) “IL-1β release without cell death” triggered by LPS+ATP. 2) “Pyroptosis” displaying IL-1β and cell death, triggered by LPS+nigericin. 3) “Cytokine uncoupled pyroptosis” presents cell death without cytokine release, triggered by nigericin. We further delineate the differential signalling involved in these distinct responses. This study also unravels the role of DNA damage-cytoskeletal reorganization in nuclear rounding and pyroptotic cell death.

Multiple studies have suggested that neutrophils resist pyroptosis and release IL-1β independent of cell death (6, 7, 12, 20). Notably, Karmakar et al., (6) showed murine neutrophils to be resistant to cell death with a short 60-90 min exposure to inflammasome activators, including nigericin and ATP, in LPS-primed cells. While in the present study, we observed increased pyroptotic cell death in neutrophils upon nigericin treatment when analyzed over a longer period (180 min). This includes hallmark features of pyroptotic cell death, including cell swelling/ballooning, pore formation, and rupture. Consistent with the previous reports, we observed IL-1β release without cell death in the case of LPS-primed ATP-treated neutrophils (6). These findings are in contrast to prior studies in macrophages, where LPS+ATP induces both IL-1β release and cell death (6, 16) and no IL-1β and LDH release observed with nigericin alone (16). This advocates differential machinery in neutrophils and macrophages, leading to distinct outcomes in response to pyroptotic stimuli including nigericin and ATP.

Our mechanistic analysis confirmed pyroptosis in LPS-primed nigericin-treated neutrophils, which displayed IL-1β release dependent on NLRP3 and caspase-1 activation. Cytokine-uncoupled pyroptosis in nigericin-treated neutrophils did not display NLRP3 and caspase-1 activation. In contrast, non-canonical caspase-11 seems critical for IL-1β from live neutrophils in case of LPS + ATP treatment. This is consistent with previous observations of Caspase-11 activation with cytosolic bacterial invasion or intracellular LPS (21, 41). Thus, LPS priming acts as a decisive factor in regulating IL-1β secretion through NLRP3 and caspase 1/11 signalling in neutrophils.

While “pyroptotic cell death” observed in nigericin-treated neutrophils, independent of LPS priming, was majorly dependent on caspase-9, caspase-7, and DNA damage in neutrophils. Moreover, LPS priming protected against DNA damage activation. Intriguingly, LPS priming supported IL-1β secretion in response to nigericin treatment, while LPS alone did not induce substantial IL-1β release in neutrophils. As expected, enhanced DNA damage with KU-55933 augmented cell death and LDH release. More surprisingly, forced DNA damage prevented NLRP3 expression and IL-1β secretion in LPS-primed nigericin-treated neutrophils and suggested the role of DNA damage in altering the cellular fate towards cytokine-uncoupled pyroptosis. The differential regulation of pyroptotic death and IL-1β secretion was further distinguished with diminished AKT-ERK signaling in nigericin-treated neutrophils that was dependent on LPS priming in neutrophils. MAPK signaling is known to regulate neutrophil survival (33, 34). Interestingly, pharmacological inhibition of MAPK signaling did not affect the pyroptotic cell death or IL-1β secretion, but reduced TNF-ɑ secretion, indicating distinct regulatory mechanisms governing the secretion of these inflammatory mediators (42).

Our data suggest that neutrophil pyroptotic death with nigericin treatment depends on GSDMD-dependent pore formation in LPS priming independent manner. While negligible pores were observed in ATP-treated cells, further suggesting a Gasdermin-D-independent pathway for IL-1β secretion from live neutrophils, as reported in studies demonstrating IL-1β secretion from viable cells, including dendritic cells and monocytes (6, 23). No uptake of small dyes, PI, and EtBR, confirmed the same. Gasdermin-D pore inhibition with necrosulfonamide blocked IL-1β and LDH release in LPS-primed nigericin-treated neutrophils, as observed earlier in macrophages (43). It is important to mention that our SEM study revealed pores of varying diameter, ranging from 20 nm to >100 nm, on neutrophil cell surfaces. These heterogeneous pores are likely driven by the collision of small pores on the cellular membrane, which is supported by the sequential permeability of DNA binding dyes - ethidium bromide (394 Da), Sytox Green (609 Da), and propidium iodide (PI, 668 Da) as per their molecular weight suggests a progressive increase in pore size (32, 44). Conversely, it is important to mention that the uniformity of pores measuring 18-23 nm is typically observed in liposomes, whereas heterogeneous oligomers were observed in cells overexpressing GSDMD-NT (45). This supports the presence of heterogeneous pores on biological membranes, which remain less described for unknown reasons. Furthermore, our study revealed mitochondrial-independent IL-1β secretion and LDH release in neutrophils. This is consistent with a previous study that demonstrated mitochondrial-independent NLRP3-caspase-1 activation, IL-1β secretion, and LDH release in LeTx-induced pyroptosis (30).

Our data also recognize nuclear rounding preceding cell swelling/ ballooning, followed by loss of plasma membrane permeability. Moreover, the perinuclear actomyosin contractile forces facilitated the process of nuclear rounding. The enhanced F-actin organization in the perinuclear region supports the putative inter-organelle communication regulating cell death and IL-1β response. Modulation of cytoskeletal remodeling via Y-27632 inhibited nuclear rounding and cell death but did not affect IL-1β secretion in neutrophils. This is consistent with our previous observation of Y-27632 independent IL-1β level in bronchoalveolar lavage (BAL) fluid neutrophils (46).

The neutrophil’s multi-lobulated nucleus is structured differently from the nucleus in most of the other cell types and has been used historically for neutrophil identification and characterization (47). Still relevance of the neutrophil nucleus remains limitedly explored in migration with putative flexibility and biomechanics (48). Deformability of the nucleus remains a puzzling question, which is not static and displays hyper-segmentation, hypolobulation, condensation, decondensation or swelling (49, 50) and compression leading to rounding, observed in this study. Recent understanding of neutrophil plasticity further supports the enigmatic nature of neutrophil nucleus, particularly under inflammatory and disease conditions. These data suggest the possibility of misinterpretation of neutrophil numbers in pathological conditions, which may be due to the presence of neutrophils with round-shaped nucleus, as we observed in the acute lung injury model. Low-density neutrophils (LDN) with less-segmented nuclei often present in systemic lupus erythematosus (SLE) and rheumatoid arthritis (RA) patients, which includes mature neutrophils with delobulation (51, 52). Moreover, Pelger-Huët anomaly (PHA), a rare genetic disorder resulting from an autosomal dominant LBR mutation, is characterized by hypo-lobulated, ovoid neutrophil nucleus (50, 53). Future studies are warranted to understand changes and crosstalk of nuclear membrane proteins, including Lamins, receptors, and cytoskeletal proteins regulating the reorganization of neutrophil nuclear structure during pyroptosis.

## CONCLUSIONS

This study demonstrates pyroptotic responses, particularly the inter-regulation of IL-1β release and pyroptotic cell death in neutrophils. Furthermore, our data unravel the cellular and molecular events underlying neutrophil pyroptotic cell death and find actomyosin signaling regulating nuclear reorganization during pyroptotic death in neutrophils. The bacteria-induced neutrophil pyroptosis exhibiting nucleus rounding both under *in-vitro* and *in-vivo* settings further validates the broad nature of this phenomenon. Human neutrophils also exhibited an association of nuclear rounding with pyroptotic cell death. Our findings illuminate the distinct mechanisms leading to the features of cell death and cytokine secretion, including DNA damage, NLRP3 inflammasome activation, nuclear rounding, and cell swelling during pyroptosis in neutrophils. The absence of LPS priming leads to the uncoupling of IL-1β and pyroptotic cell death based on a lack of NLRP3 and caspase-1 activation in neutrophils, which can be regulated by distinct cues and conditions (54). Intriguingly, forced DNA damage caused death, but surprisingly blocked IL-1β secretion. It will be interesting to further understand the immunogenic or immunomodulatory consequences of these distinct decisions. Together, further understanding and targeting these distinct pathways may provide specific outcomes and may have broad biological significance in various pathological conditions.

## MATERIAL AND METHODS

### Reagents and antibodies

All reagents and antibodies used in the study are listed in the table provided in the supplementary file.

### Animals

The C57BL/6 mice (male; eight to twelve weeks old) were used for the experiments. Mice were bred and kept in the animal house at CSIR-CDRI, and all the experiments were performed as per the IAEC guidelines.

### Neutrophil isolation

Murine neutrophils were isolated from mouse bone marrow using Percoll density gradient as described previously (46). Briefly, bone marrow cells were flushed out with Hank’s balanced salt solution (HBSS) containing 0.1% bovine serum albumin (BSA). Cells were layered over a discontinuous Percoll gradient (52, 62, and 70%), and centrifuged (750 g, 25 min, room temperature (RT). The interface between the 62 and 70%-layers containing neutrophils was harvested and washed with HBSS containing 0.1% BSA. RBCs were removed using Histopaque-1119. Neutrophils were obtained with >95% purity and were morphologically mature. Human neutrophils were isolated from human blood using a Percoll density gradient as described previously (55).

### Cell culture and treatment

Neutrophils were cultured in RPMI-1640 supplemented with 10% heat-inactivated fetal bovine serum (FBS, Gibco) containing 2% penicillin-streptomycin (PS) antibiotics (Invitrogen) at a density of 1 × 10^6^ cells/ml at 37°C in a 5% CO2 incubator. Neutrophils were primed or not primed with 500 ng/mL LPS (*Escherichia coli O111:B4*) for 3 h in RPMI supplemented with 5% FBS and 2% penicillin-streptomycin antibiotics at 37°C in a 5% CO2 incubator. Following priming, cells were treated with inflammasome activators including nigericin 100 nM to 10 µM or ATP 5 mM (Both obtained from Cayman chemicals) for 3 h (6). ATP was dissolved in HBSS titrated to pH 7.2 with 5 M NaOH solution.

In case of testing different pathways, cells were pre-treated with different target inhibitors including Y27632 (30 µM, ROCK inhibitor), Necrosulfonamide (30 µM, GSDMD inhibitor), Q**-**VD**-**OPh (30 µM, Pan Caspase inhibitor), VX765 (30 µM, Caspase-1 inhibitor), MK2206 (1 µM, Akt inhibitor), MCC950 (30 µM, NLRP3 inflammasome inhibitor), KU-55933 (10 µM, ATM kinase inhibitor), Rotenone (10 µM, electron transport chain (ETC) - Complex 1 inhibitor), FCCP (10 µM, Protonophore inhibitor), Oligomycin (10 µM, ATP synthase inhibitor), SB203580 (1 µM, p38 MAPK inhibitor), Wortmannin (5 µM, PI3K inhibitor) (all obtained from Cayman chemicals), Caspase-9 inhibitor (50 µM, Sigma Aldrich), Antimycin (10 µM, ETC Complex IV, Sigma Aldrich) 30 min before nigericin addition.

### Peritoneal macrophages isolation and culture

For peritoneal macrophages, 5 mL of cold HBSS was injected into mice peritoneal cavity. Cells were pelleted at 500 g and resuspended in RPMI supplemented with 10% heat-inactivated FBS containing 2% P/S and allowed to adhere in a 24-well plate at a density of 1 × 10^6^ cells for 2 h. Cells were then washed with HBSS to remove non-adherent cells and cultured in RPMI supplemented with 10% heat-inactivated fetal bovine serum and used for the assay after 12 h.

### Cell membrane permeability analysis using Sytox Green or Propidium iodide

Peritoneal macrophages and neutrophils were labelled for 20 min with Hoechst-33342 (1 μg/mL, Sigma Aldrich) dye to stain nucleus of all the cells and Sytox Green (100 nM, Invitrogen), a cell-impermeable DNA binding dye to visualize cell death as described previously (55). Cells were primed or not primed with LPS from *Escherichia coli O111:B4* (500 ng/mL, Sigma Aldrich) for 3 h and then stimulated with nigericin 10 µM and ATP 5 mM for 1 or 3 h at 37°C. Cell death was imaged with a 20x objective at HCS Cellomics™ (Thermo-Fisher Scientific ArrayScan™ VTI) or DMI6000 fluorescence microscope as described previously.

For time-dependent Sytox Green uptake, 4 x 10^4^ neutrophils were labeled with Sytox Green (100 nM) in a 96-well clear-bottom black plate. After two to three baseline readings, primed or not primed neutrophils were treated with nigericin 10 µM and ATP 5 mM and fluorescence readings were recorded every 15 min up to 4.5 h using a Tecan Infinite^®^ 200 PRO plate reader (6). For PI uptake analysis by flow cytometry, neutrophils were primed with LPS and then stimulated with nigericin 10 µM or ATP 5 mM. After stimulation, cells were labelled with PI (1 µg/mL) and acquired using FACS Lyric (BD Biosciences).

### Live cell imaging

1 × 10^6^ neutrophils were pre-labelled with Calcein AM (2 µM, Invitrogen), JC-1 (500 ng/mL, Cayman Chemicals), Hoechst-33342 (1 µg/mL, Sigma Aldrich), DAPI (100 ng/mL, Sigma Aldrich), Sytox Green (100 nM, Invitrogen) or PI (1 µg/mL, Sigma Aldrich) in different combinations as mentioned in figure legends. Cells were labelled for 20 min and were seeded in a glass-bottom plate, and images of the Differential interference contrast (DIC), fluorescence channels were acquired every 10 min after nigericin addition for 3 h at 37°C at 40×/1.3 NA objective using a Leica DMI6000 fluorescence microscope. Subsequently, images were processed, and videos were generated using Leica software. Fluorescence intensity quantifications were performed using ImageJ software on cells undergoing nuclear rounding in response to pyroptotic stimuli.

### LDH cytotoxicity assay

2 x 10^5^ neutrophils primed or not primed with LPS (500 ng/mL) for 3 h were stimulated with inflammasome activators, including nigericin 10 µM or ATP 5 mM, for 5 h in RPMI containing 10% FBS at 37°C. For inhibitor screening, cells were pre-incubated with different inhibitors for 30 min before nigericin addition. LDH released was determined using the Cytotoxicity detection kit (MERCK) as per the manufacturer’s protocol. LDH release was calculated using the formula (LDH sample – LDH negative control)/ (LDH 100% lysis – LDH negative control) × 100). Media only was used as a negative control, while 100% lysis control was prepared by adding Triton X-100 to a final concentration of 1%.

### Flow cytometry time kinetics

2 x 10^6^ neutrophils were primed with LPS (500 ng/mL) in RPMI containing 5% FBS for 3 h at 37°C. After priming, cells were stained with Hoechst-33342 (1 µg/mL, Sigma Aldrich) and Sytox Green (100 nM, Invitrogen) for 20 minutes at 37°C. Cells stimulated with and without nigericin 50 µM were acquired for 1 h in time mode using FACS Aria (BD Biosciences) (31, 56).

### Enzyme-linked immunosorbent assay (ELISA) for cytokine release

2 x 10^6^ neutrophils primed or not primed with LPS (500 ng/mL), or zymosan (1 µg/mL) for 3 h were treated with inflammasome activators including nigericin 10 µM or ATP 5 mM for 1 h and incubated at 37°C in RPMI containing 5% FBS. For screening studies, cells were pre-treated with different inhibitors for 30 min before nigericin addition. After treatments, cells were centrifuged at 12,000 g for 15 min at 4°C and supernatant was collected and assayed for IL-1β, TNF-ɑ using ELISA as per manufacturer instructions (R&D Biosystems).

### Bacterial infection

*Escherichia coli* (strain 25922) colonies, grown on an agar plate, were inoculated in Luria-Bertani (LB) broth medium overnight at 37°C with shaking at 200 r.p.m. Mid-logarithmic phase bacteria, were obtained after re-inoculation into fresh LB medium grown for 2 h till an optical density (600 nm) of ∼ 0.4 was acquired. Bacteria were washed with HBSS and opsonized with murine serum for 30 min at 37°C (8). For Heat inactivation, bacteria were heated at 60⁰C for 1h in a dry bath. For LDH or IL-1β release analysis, 0.5 x 10^6^ murine neutrophils were infected with *E.coli* at a multiplicity of infection (MOI) of 10 in RPMI containing 10% FBS for 3 h. After treatment, cells were centrifuged at 2,000 g for 10 min and supernatant was analyzed for LDH or IL-1β release as described above. For nuclear fractionation experiments, 4 x 10^6^ neutrophils/mL were cultured with *E. coli* at MOI of neutrophils to bacteria (1:10) for 15 min and washed with HBSS containing 0.1% BSA at 200 g for 10 min to remove extracellular bacteria. *E. coli* challenged neutrophils were further cultured for 1.5 h in RPMI containing 10% FBS to probe nuclear changes using Western blot analysis (8).

### Cytosolic and Nuclear fractionation

4 x 10^6^ murine neutrophils were cultured in 1 mL of RPMI containing 10% FBS with nigericin 10 µM or *E.coli* at a MOI of 10 for 1.5 h. After nigericin treatment, cells were pelleted at 12,000 g for 10 min. While, in case of nuclear cytoplasmic fractionation from *E. coli-treated* neutrophils, cells were pelleted at 220 g for 5 min to prevent any actin inclusion from bacteria in the nuclear or cytoplasmic fraction. Cell pellets were resuspended in 150 µL of hypotonic lysis buffer [10 mM Hepes (pH 8.0), 10 mM KCl, 3 mM NaCl, 3 mM MgCl2, 1 mM EDTA, 1 mM EGTA, 2 mM dithiothreitol (DTT), protease inhibitor cocktail (Sigma Aldrich)] for 15 min on ice. After incubation 7.5 μL of 10% Nonidet P-40 was added, and the cells were gently vortexed, immediately centrifuged at 500 g for 10 min at 4°C. The supernatant was collected and designated as cytoplasmic extracts. Pelleted nuclei were washed with 200 µL hypotonic lysis buffer containing protease inhibitor cocktail, resuspended in 100 µL of RIPA lysis buffer, sonicated, centrifuged (12,000 g, 15 min, 4°C) and the resulting supernatants - nuclear extracts were used to check for the expression of β-actin, Histone 3B (all from Cell Signaling Technology, Boston, MA). Cytosolic and nuclear fractionation was confirmed using ɑ-tubulin or Histone 3B respectively (57).

### Western Blotting

2 x10^6^ neutrophils were primed with LPS (500ng/mL) for 3 h and stimulated with nigericin or ATP for 1 or 3 h at 37°C, as indicated in the figures. Cells were lysed using RIPA lysis buffer (150 mM NaCl, 50 mM Tris-HCl, 1% Triton X-100, 0.1% SDS) or Caspase lysis buffer to check Caspase activation (50 mM HEPES, 100 mM NaCl, 1mM EDTA, 0.5% Triton-X, 10% Sucrose, 4% Glycerol, 5 mM DTT, 0.1% CHAPS) supplemented with protease inhibitor cocktail. Cell lysate containing an equal amount of protein was separated on SDS-PAGE and analyzed for the expression of NLRP3, p γH2AX (Ser 139), p-Akt (Ser 473), p-ERK1/2 or Cleaved Caspase-7, Caspase-9, Caspase 11 and β-actin (all obtained from Cell Signaling Technology, Boston, MA, and used at a dilution of 1:1000).

For analysis of secreted proteins/cytokines after stimulation with nigericin or ATP, cells were centrifuged at 12,000 g for 10 min at room temperature, the supernatant was mixed with methanol and chloroform at a ratio of 1:4. The interface layer containing precipitated protein was collected, washed with methanol and air dried. Pellet was resuspended in 2X loading dye. 15 µg of total cell lysate and precipitated supernatant of 1 million neutrophils was mixed and analysed using western blotting for cleaved caspase-1 (1:1000), NLRP3 (1:1000) or mature IL-1β (1:1000) (all obtained from Cell Signaling Technology, Boston, MA). Western analysis was performed using Image J software and reported as % of maximum calculated as measured band intensity / maximum band intensity × 100. Band intensity represented the value after actin normalization (56).

### Scanning electron microscopy

1 x 10^6^ neutrophils were cultured in RPMI containing 10% FBS in 24 well plate on 10 mm coverslip coated with 0.01% poly-L-lysine with or without LPS priming (500 ng/mL) for 3 h and then stimulated with nigericin or ATP for 1 h at 37°C. Cells were fixed overnight with 2.5% glutaraldehyde prepared in phosphate buffer (pH 7.4) at 4°C. Cells were washed with PBS followed by the addition of 1% Osmium tetraoxide in phosphate buffer saline and incubated for 1 h. After washing, cells were dehydrated in increasing concentrations of ethanol, followed by critical point drying in critical point dryer and sputter coated with Au: Pd (80:20) and visualized using FEI Quanta 250 Scanning electron microscope at 20Kv, using SE detector (58). Pore sizes were calculated using Image J software (45).

### Immunofluorescence staining

30,000 neutrophils with or without LPS priming (500ng/mL), were stimulated with 10 µM nigericin or 5 mM ATP and seeded on a multi-chamber slide for 3 h in a humidified chamber at 37°C. After incubation, cells were fixed with 2% paraformaldehyde (PFA) for 15 min. Cells were permeabilized with 0.1% Triton X, blocked with 2% BSA, and stained overnight with primary antibodies as follows-8-OHdG (1:1000) (Obtained from Merck), IL-1β (1:100) (Invitrogen) or pMLC (1:100) (Myosin light chain; Cell Signaling Technology, Boston, MA) overnight at 4°C. Cells were then washed and stained with secondary antibody anti-goat Alexa fluor 488 or anti-rabbit Alexa fluor 488 for 2 h at RT (1:500, Invitrogen) and mounted in DAPI-containing media (Invitrogen) (59).

For F-actin staining, cells were fixed with 4% PFA for 15 min. F-actin was stained using rhodamine-phalloidin (1:100) for 2 h at room temperature and mounted in DAPI-containing media (Invitrogen). Images were acquired using a Leica DMI6000 fluorescence microscope at 40×/1.3 NA objective or an Olympus confocal microscope at 60x objective. The percentage of cells with segmented, intermediate and round nucleus after nigericin treatment with or without different inhibitors was enumerated in ten to fifteen non-overlapping fields of view, with a minimum of 50 cells/ field and represents data from a minimum of 3 separate sets of experiments.

### Mitochondrial damage detection

2 x 10^5^ neutrophils were primed or not primed with LPS (500 ng/mL) for 1.5 h and then stimulated with nigericin 10 μM or ATP 5 mM for 2 h at 37°C. For inhibitor treatment, cells were pre-treated for 30 min and then stimulated with nigericin or ATP. Mitochondrial mass and membrane potential measurements were made using 50 nM of MitoTracker Green and MitoTracker Red CMXRos dye (Both obtained from Invitrogen), stained for 30 min at 37⁰C. Mitochondrial ROS was detected using MitoSOX (Invitrogen) at 2 µM for 30 min at 37⁰C (29).

### *E. coli* and LPS-induced acute Lung Injury Model

The C57BL/6 mice were challenged with 0.5 million cfu of heat-inactivated *Escherichia coli* (strain 25922) or LPS (1.5mg/kg) suspended in HBSS by intratracheal instillation after ketamine and xylazine anesthesia. Heat-inactivated *E. coli* was used to just trigger neutrophil influx and to avoid *E.coli* growth and infection *in-vivo.* Bronchoalveolar Lavage Fluid (BALF) was collected after 24 h of *E.Coli* or LPS challenge. BALF was centrifuged at 500 g for 10 min and used for further assays (46).

For Giemsa staining, BALF cells (2 x10^4^ cells) collected from the lung were centrifuged at 500 g for 5 min on slides. Cells were fixed with a May-Grunwald stain for 20 min. Slides were then washed, dried, and mounted with DPX mounting medium. Images were captured and analyzed using a Nikon Eclipse TS2 microscope with a 60× objective.

For elastase staining, BALF cells were fixed using 2% PFA for 15 min and stained with a primary antibody (1:100) for neutrophil elastase for 2 h. Cells were then washed and stained with secondary antibody anti-rabbit Alexa fluor 568 for 2 h at RT (1:500, Invitrogen) and mounted in DAPI-containing media (Invitrogen). Images were acquired using an Olympus confocal microscope at a 60x objective. The percentage of cells with a segmented and round nucleus containing neutrophil elastase was enumerated in 100 cells and represents data from 3 independent experiments.

### Statistical analysis

All the experiments were performed at least three times. An unpaired Student’s *t-*test (normally distributed), One-way or Two-way ANOVA were performed as statistics using Prism 8 software (GraphPad) to compare experimental groups as mentioned in the figure legends unless specified. Data are mean ± SEM. The p-value of * P *<* 0.05; ** P *<* 0.01; and *** P *<* 0.001, **** P *<* 0.0001 were considered as significant.

## Supporting information

supplementary file Singhal etal

## ACKNOWLEDGEMENTS

We thank Mr. Anil Kumar Verma at Confocal/intravital facility for his help with confocal experiments at CSIR-CDRI. We thank Dr. CP Pandey for his support at fluorescence imaging facility at the Department of Pharmacology at CSIR-CDRI. We thank Mr. AL Vishwakarma at the Flow Cytometry Core for his help with flow cytometry experiments. We thank the Director CSIR-CDRI for her support during the study. We acknowledge the financial support of the early career grant from DST-SERB ECR/2022/001339, CSIR-FBR070302, BT/CS0075/06/22. MD is supported by JBR/2020/000034. This is CDRI communication number 184/2024/SK.

## AUTHOR CONTRIBUTIONS

A.S performed, analyzed experiments, interpreted data, prepared figures and wrote the manuscript. N.A, P.S.M, and R.G performed and analyzed specific experiments. N.C, and K.M performed, interpreted and analyzed the scanning electron microscopy experiments and data analysis. T.C and M.D. provided critical inputs in experimental design. S.K. designed, interpreted experiments and wrote the manuscript.

## DECLARATION OF GENERATIVE AI AND AI-ASSISTED TECHNOLOGIES

During the finalization of this work, the author(s) used the Grammarly generative AI tool in order to improve language and readability. After using this tool, the author(s) reviewed and edited the content as needed and take(s) full responsibility for the content of the publication.

