## supplementary file Singhal etal for "Inflammasomes and DNA Damage Orchestrate Divergent Pyroptotic Fates in Neutrophils"

### Supplementary figure and legends

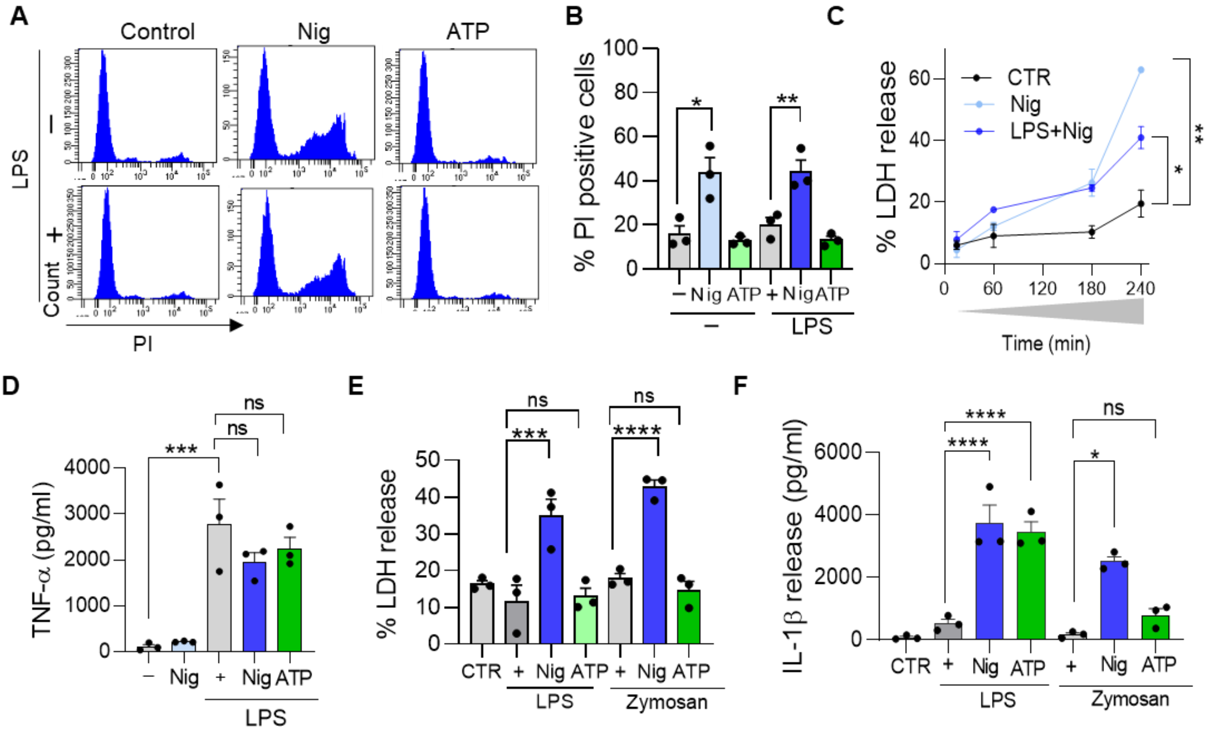


**Supplementary Fig 1: Additional analysis of Cell death and cytokine release.**

**(A)** Representative histogram for PI with nigericin or ATP treatment in the presence and absence of LPS priming in neutrophils.

**(B)** Quantification of % PI-positive cells with nigericin or ATP treatment in the presence and absence of LPS priming in neutrophils (*P <0.05, **P <0.01, n = 3 independent experiments, analyzed using One-way ANOVA).

**(C)** Time-dependent analysis for LDH with nigericin in the presence and absence of LPS priming (*P <0.05, **P <0.01, n = 3 independent experiments, analyzed using unpaired t test).

**(D)** TNF-α (pg/ml) release with nigericin and ATP with and without LPS priming in neutrophils. (***P <0.001, ns - not significant, n=3 independent experiments, analyzed using One-way ANOVA).

**(E)** LDH release with nigericin and ATP in LPS and Zymosan primed neutrophils (***P <0.001, ns - not significant, n=3 independent experiments, analyzed using One-way ANOVA).

**(F)** IL-1β release (pg/ml) with nigericin and ATP in LPS and Zymosan primed neutrophils (****P <0.0001, ns - not significant, n=3 independent experiments, analyzed using One-way ANOVA).

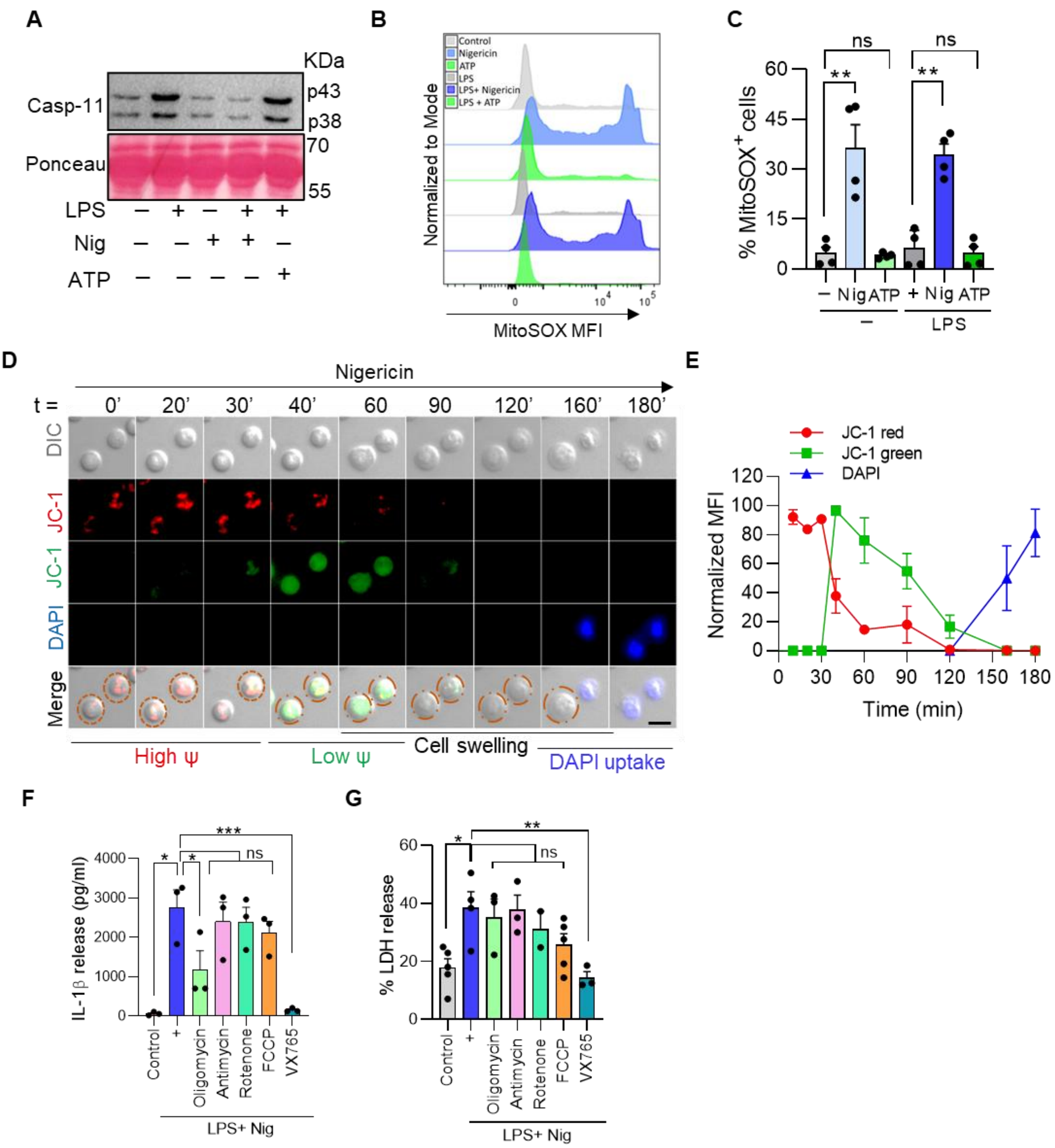


**Supplementary Fig 2: Effect of Caspase-1 and NLRP3 inflammasome inhibitor on IL-1β release and cell death.**

**(A)** Western blot image for caspase-11 expression in nigericin and ATP-treated neutrophils with and without LPS priming.

**(B)** Representative histogram shows MitoSOX Mean fluorescence intensity with and without LPS priming in nigericin and ATP-treated neutrophils.

**(C)** % MitoSOX positive cells in nigericin and ATP-treated neutrophils with and without LPS priming (**P <0.01, n = 4 independent experiments, analyzed using One-way ANOVA).

**(D)** Representative image of time-dependent changes in mitochondrial membrane potential (high (red) /low (green)) labelled with JC-1 and DAPI (blue) in nigericin-treated neutrophils (Scale bar, 10 μm).

**(E)** Quantification of time-dependent changes in mitochondrial membrane potential (high (red) /low (green)) labelled with JC-1 and DAPI (blue) in response to nigericin.

**(F)** IL-1β release in LPS + nigericin-challenged neutrophils in the presence or absence of mitochondrial modulators: FCCP (Protonophore), oligomycin (ATP synthase inhibitor), rotenone (Electron transport chain (ETC) - Complex I inhibitor), and antimycin (ETC - Complex IV inhibitor) (*P<0.05, ***P <0.001, n = 3 independent experiments, analyzed using One-way ANOVA).

**(G)** LDH release in LPS + nigericin-challenged neutrophils in the presence or absence of mitochondrial modulators (*P<0.05, **P <0.01, n = 3 independent experiments, analyzed using One-way ANOVA).

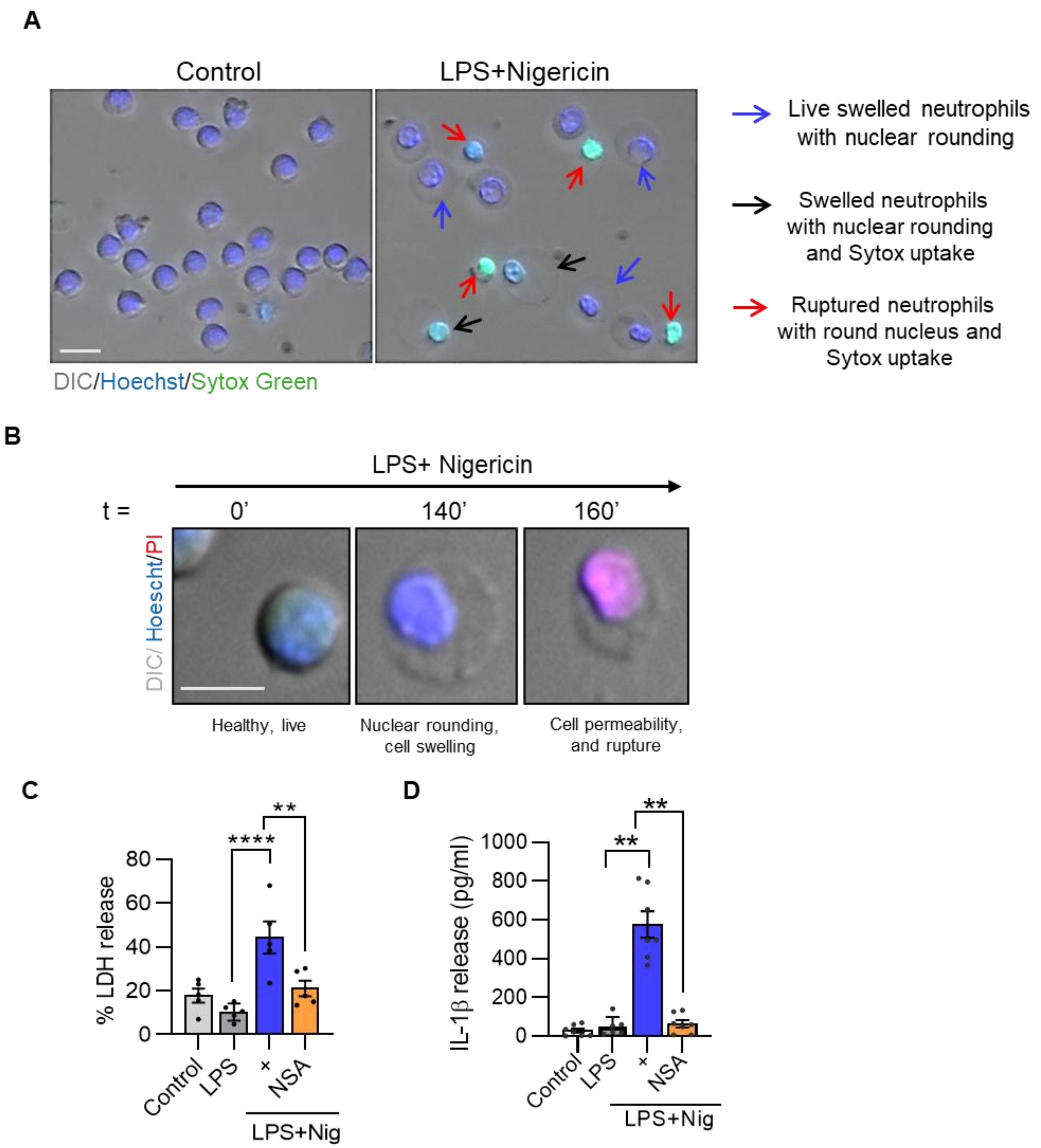


**Supplementary Fig 3: Cellular events during pyroptosis in neutrophils.**

**(A)** Representative image showing DIC (gray), Hoechst (blue), SYTOX Green (green) and distinct stages observed in LPS+nigericin-treated neutrophils (n = 3 independent experiments with a minimum of 100 cells per experiment) (Scale bar, 20 μm). The blue arrow indicates cells with swelling, round nuclei with intact membranes. The black arrow indicates cells showing swelling, SYTOX green-positive round nuclei indicating compromised plasma membrane integrity. The red arrow indicates cells that show ruptured cells with SYTOX green-positive nuclei, indicating loss of cellular content.

**(B)** Representative image showing sequence of events in a single cell traced after LPS+nigericin addition using DIC (grey), Hoechst (blue) and PI (red) (n = 2 independent experiments, with a minimum of 50 cells analyzed per experiment) (Scale bar, 20 μm).

**(C)** % LDH release in the presence of necrosulfonamide (NSA - Gasdermin D inhibitor - 30μM) in LPS+nigericin treated neutrophils (**P <0.01, ****P <0.0001, n = 4 independent experiments, analyzed using One-way ANOVA).

**(D)** IL-1β release in the presence of necrosulfonamide (NSA - Gasdermin D inhibitor - 30μM) in LPS+nigericin treated neutrophils (**P <0.01, ****P <0.0001, n = 7 independent experiments, analyzed using One-way ANOVA).

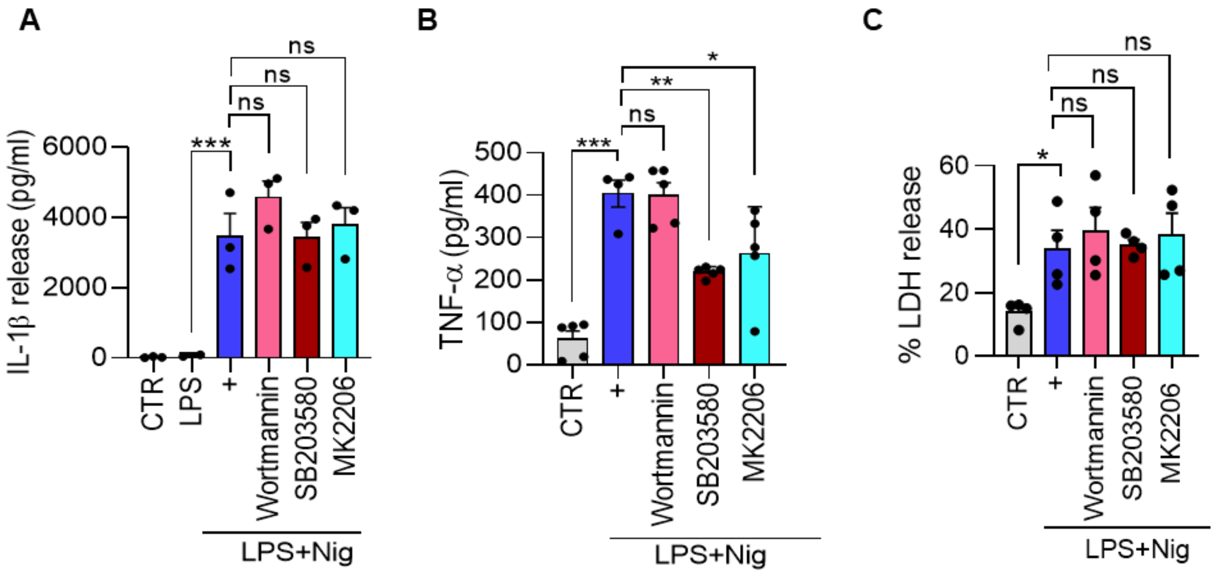


**Supplementary Fig 4: Distinct regulation of TNFα and IL-1β release via MAPK signalling.**

**(A)** Effect of different inhibitors for MAPK signalling – Wortmannin (PI3 Kinase), SB203580 (p38 MAPK) and MK2206 (Akt inhibitor) on IL-1β release (***P <0.001, ns - not significant, n = 3 independent experiments, analyzed using One-way ANOVA).

**(B)** Effect of Wortmannin, SB203580 and MK2206, on TNF-α secretion release in LPS + nigericin treated neutrophils (***P <0.001, n = 4 independent experiments, analyzed using One-way ANOVA).

**(C)** Effect of Wortmannin, SB203580 and MK2206 on LDH release in LPS + nigericin-treated neutrophils (*P <0.05, n = 4 independent experiments, analyzed using One-way ANOVA).

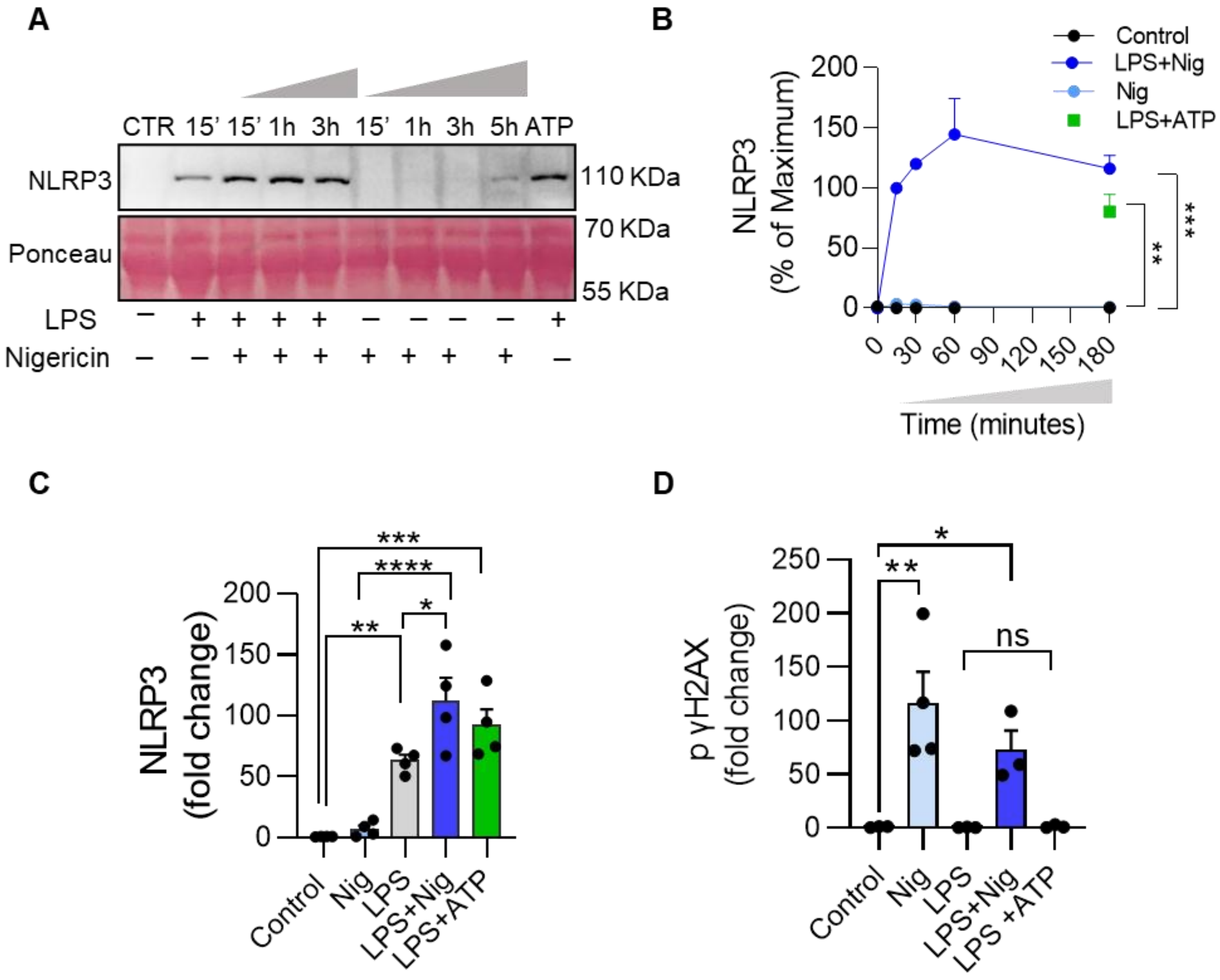


**Supplementary Fig 5: γH2AX and NLRP3 expression under different conditions.**

**(A)** Western blot image for time-dependent NLRP3 expression with nigericin treatment in the presence and absence of LPS priming.

**(B)** Quantification of time-dependent NLRP3 expression with nigericin or ATP treatment in the presence and absence of LPS priming (**P <0.01, ***P <0.001, n = 3 independent experiments, analyzed using One-way ANOVA).

**(C)** Quantification of NLRP3 fold change with nigericin or ATP treatment in the presence and absence of LPS priming (*P <0.05, **P <0.01, ***P <0.001, ****P <0.0001, n = 3 independent experiments, analyzed using One-way ANOVA).

**(D)** Quantification of p γH2AX with nigericin and ATP in the presence and absence of LPS priming (*P <0.05, **P <0.01, n = 4 independent experiments, analyzed using One-way ANOVA).

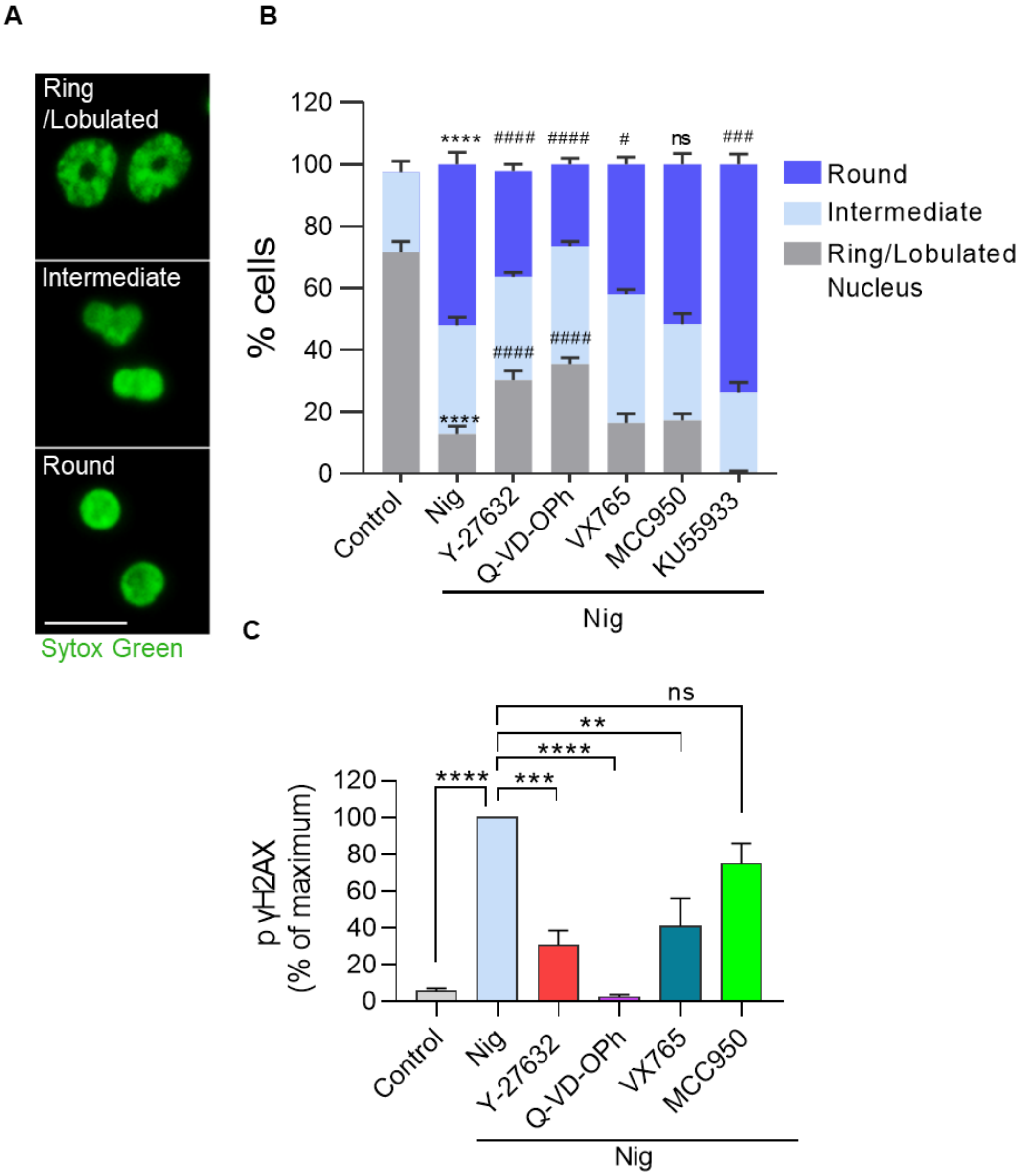


**Supplementary Fig 6: Analysis of different nuclear phenotypes/morphology observed in nigericin-stimulated neutrophils.**

**(A)** Representative image showing distinct nuclear phenotypes observed in nigericin-treated neutrophils stained with SYTOX Green (green) (Scale bar, 10 µm).

**(B)** Relative quantification showing the effect of different inhibitors on nuclear morphology in nigericin-stimulated neutrophils. (#P <0.05, ****P <0.0001, ####P <0.001, n = 4 independent experiments, with a minimum of 100 cells per experiment, analyzed using Two-way ANOVA).

**(C)** Quantifications showing the effect of different inhibitors on p γH2AX expression in nigericin-treated PMNs (**P <0.01, ***P <0.001, ****P <0.0001, n = 3 independent experiments, analyzed using One-way ANOVA).

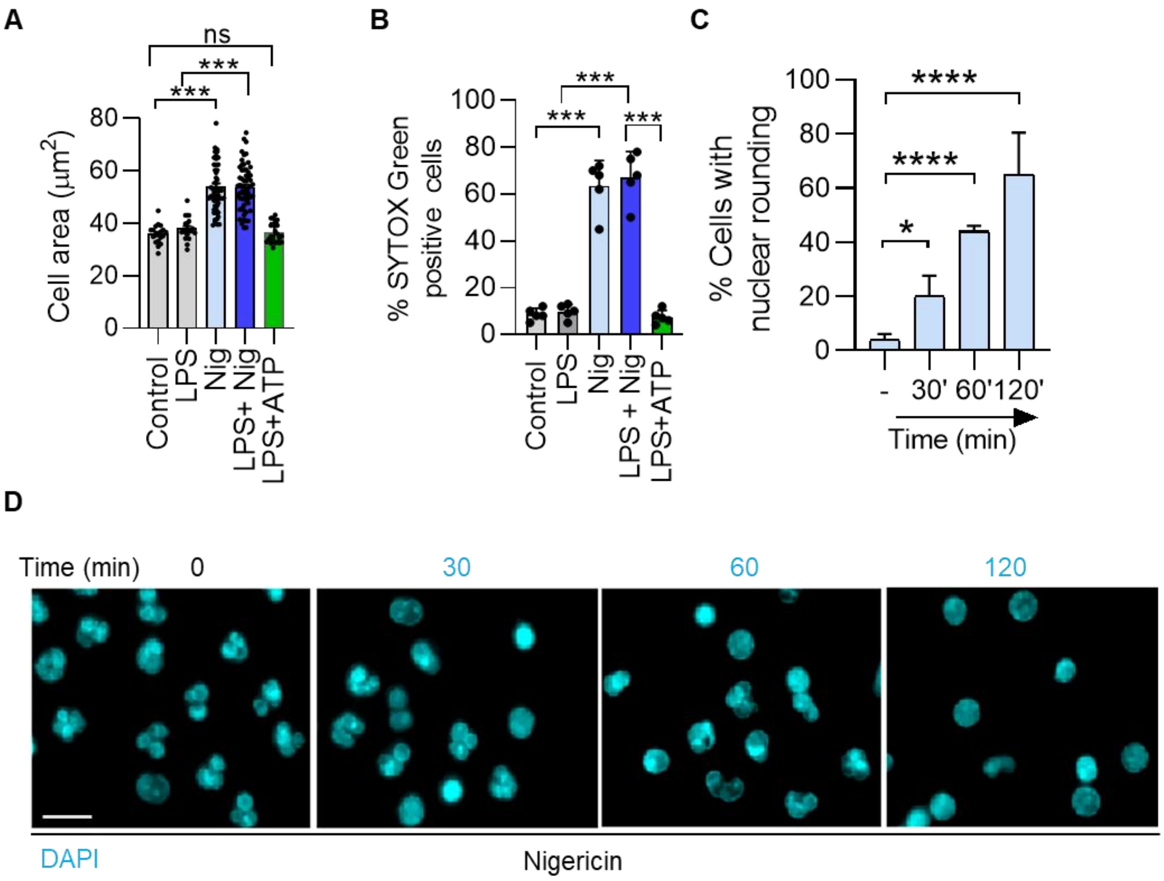


**Supplementary Fig 7: Human neutrophils exhibit similar nuclear rounding, cell swelling and cell death in different conditions.**

**(A)** Cell area (μm$^2$) of neutrophils after nigericin and ATP treatments in the presence and absence of LPS priming (***P <0.001, n = 3-4 independent experiments, analyzed using One-way ANOVA).

**(B)** % SYTOX Green positive cells after nigericin and ATP treatments in the presence and absence of LPS priming (***P <0.001, n = 4 independent experiments, analyzed using One-way ANOVA).

**(C)** Percentage of cells with nuclear rounding after nigericin and ATP treatments in the presence and absence of LPS priming (*P <0.05, ****P <0.0001, n = 3 independent experiments, analyzed using One-way ANOVA).

**(D)** Representative image showing time-dependent nuclear changes after nigericin addition, stained with DAPI (Cyan) (Scale bar, 10 μm).

Table 1. Resource Table.

| REAGENT or RESOURCE | SOURCE | IDENTIFIER |
|---|---|---|
| **Antibodies** | | |
| **8-Hydroxy-2′-deoxyguanosine (8-oHdG)** | Merck | Cat#AB5830 |
| **Anti α-Tubulin antibody** | DHSB | Cat#12G10 |
| **Anti Histone H3 antibody** | Cell Signalling Technology | Cat#4499 |
| **Anti-mouse Horseradish peroxidase** | Cell Signalling Technology | Cat# 4370S |
| **Anti-rabbit Horseradish peroxidase** | Cell Signalling Technology | Cat#7074S |
| **Anti-rat Horseradish peroxidase** | Cell Signalling Technology | Cat#7077 |
| **Caspase 11 antibody** | Cell Signalling Technology | Cat#17D9 |
| **Caspase 9 antibody** | Cell Signalling Technology | Cat#9508S |
| **Cleaved Caspase-1 (Asp296) antibody (E2G2I)** | Cell Signalling Technology | Cat#89332 |
| **Cleaved Caspase-7 (Asp198) antibody (D6H1)** | Cell Signalling Technology | Cat#8438 |
| **Cleaved-IL-1β (Asp117) antibody (E7V2A)** | Cell Signalling Technology | Cat# 63124 |
| **Donkey anti mouse IgG Alexa Fluor 488** | Invitrogen | Cat#A21202 |
| **Donkey anti rabbit IgG Alexa Fluor 488** | Invitrogen | Cat# A21206 |
| **Donkey anti rabbit IgG Alexa Fluor 568** | Invitrogen | Cat# A10042 |
| **IL-1β Recombinant Rabbit monoclonal antibody** | Merck | Cat#MA5-47038 |
| **Neutrophil elastase antibody** | Calbiochem | Cat#481001 |
| **NLRP3 (D4D8T) antibody** | Cell Signalling Technology | Cat#15101S |
| **p γH2AX (Ser 139) antibody** | Cell Signalling Technology | Cat#2577S |
| **Phosho-Myosin light chain II antibody** | Cell Signalling Technology | Cat#7074P2 |
| **Phospho-Akt antibody** | Cell Signalling Technology | Cat#4060S |
| **Phospho-ERK1/2 antibody** | Cell Signalling Technology | Cat#7074S |
| **β actin antibody** | Sigma Aldrich | Cat#A3854 |
| **Dyes, inhibitor, other reagents** | | |
| **Ammonium persulfate** | Sigma Aldrich | Cat#A3678 |
| **Antimycin** | Sigma Aldrich | Cat#A-8674 |
| **Acrylamide** | Sigma Aldrich | Cat#A8887 |
| **ATP sodium salt** | Cayman Chemicals | Cat#14498 |
| **Bicinchoninic acid (BCA)** | G Biosciences | Cat#786-846 |
| **Bisacrylamide** | Sigma Aldrich | Cat#146072 |
| **Bovine serum albumin fraction V (BSA)** | SRL chem | Cat#83803 |
| **Calcien AM** | Invitrogen | Cat#C-1430 |
| **Caspase-9 Inhibitor II, Cell-Permeable** | Sigma Aldrich | Cat#218776 |
| **Chloroform** | SRL chem | Cat#96764 |
| **CHAPS** | SRL chem | Cat#21420 |
| **Dithiothreitol (DTT)** | Roche | Cat#10708984001 |
| **Ethylenediaminetetraacetic acid (EDTA)** | Sigma Aldrich | Cat#E-6511 |
| **Ethyleneglycol-bis(β-aminoethyl)-N,N,N',N'-tetraacetic Acid (EGTA)** | ACROS ORGANICS | Cat#67-42-5 |

| REAGENT or RESOURCE | SOURCE | IDENTIFIER |
|---|---|---|
| **FCCP** | Cayman Chemicals | Cat#15218 |
| **Fetal bovine serum (FBS)** | GIBCO | Cat#10270106 |
| **Fluoromount- Mounting Medium, with DAPI** | Thermo scientific | Cat# 00-4959-52 |
| **Glutaraldehyle** | Sigma Aldrich | Cat#G5882 |
| **Glycerol** | RANKEM | Cat#G0020 |
| **HEPES** | Sigma Aldrich | Cat#15630-080 |
| **Histopaque 1119** | Sigma Aldrich | Cat#11191 |
| **10X HBSS** | Gibco | Cat# 14185052 |
| **Ionomycin** | Cayman Chemicals | Cat#10004974 |
| **4-iodophenylboronic acid (4IPBA)** | Sigma Aldrich | Cat#471933 |
| **Potassium chloride (KCl)** | Sigma Aldrich | Cat#P9333 |
| **KU-55933** | Cayman Chemicals | Cat#16336 |
| **LPS from Escherichia coli O111:B4** | Sigma Aldrich | Cat#L2630 |
| **Luminol** | | |
| **MCC950** | Cayman Chemicals | Cat#17510 |
| **Methanol** | CDH | Cat#67-56-1 |
| **Magnesium Chloride ($MgCl_2$)** | SRL chem | Cat#69396 |
| **May Grunwald** | | |
| **MK2206** | Cayman Chemicals | Cat#11593 |
| **N, N, N′, N′-Tetramethylethylenediamine (TEMED)** | Sigma Aldrich | Cat#1107320100 |
| **Nigericin sodium salt** | Cayman Chemicals | Cat#11437 |
| **Necrosulfonamide** | Cayman Chemicals | Cat#20844 |
| **3-color prestained protein ladder** | Puregene | PG-PMT2922 |
| **Oligomycin** | Cayman Chemicals | Cat#11342 |
| **Nonidet P-40** | Sigma Aldrich | Cat#74385 |
| **Protease Inhibitor Cocktail** | Sigma Aldrich | Cat#P8340 |
| **Osmium tetraoxide** | Sigma Aldrich | Cat#201030 |
| **Paraformaldehyde** | Sigma Aldrich | Cat#P6178 |
| **Penicillin-streptomycin (P/S) glutamine** | Gibco | Cat# 1881463 |
| **Percoll** | Cytiva | Cat#17089101 |
| **PMA** | Sigma Aldrich | Cat#P8139 |
| **Poly-L-lysine** | Sigma Aldrich | Cat#P8920 |
| **Ponceau S** | Sigma Aldrich | Cat#P-3504 |
| **Q-VD-OPh** | Cayman Chemicals | Cat#15260 |
| **Rhodamine-labeled phalloidin** | Invitrogen | Cat#R415 |
| **Rotenone** | Sigma Aldrich | Cat#R8875 |
| **RPMI-1640 media** | Lonza | Cat#12-702F |
| **SB203580** | Cayman Chemicals | Cat#13067 |
| **Sodium Chloride (NaCl)** | Molychem | Cat#25620 |
| **Sodium dodecyl sulfate (SDS)** | Sigma Aldrich | Cat#436143 |
| **Staurosporine** | Cayman Chemicals | Cat#81590 |
| **Sucrose** | CDH | Cat#987380 |
| **Sytox Green** | Invitrogen | Cat#S7020 |
| **Tris** | Sigma Aldrich | Cat#252859 |

| REAGENT or RESOURCE | SOURCE | IDENTIFIER |
|---|---|---|
| **Triton X-100** | Sigma Aldrich | Cat#T8787 |
| **Tween 20** | Sigma Aldrich | Cat#28599 |
| **VX765** | Cayman Chemicals | Cat#28825 |
| **Y-27632** | Cayman Chemicals | Cat#1005583 |
| **Zymosan A** | Cayman Chemicals | Cat#21175 |
| **PVDF membrane** | Millipore | IPVH00010 |
| **Hoechst- 33342** | Sigma Aldrich | Cat#B2261 |
| **Propidium iodide** | Sigma Aldrich | Cat#P4170 |
| **DAPI** | Sigma | Cat#D9542 |
| **JC-1** | Cayman Chemicals | Cat#15003 |
| **MitoTracker Green FM** | Invitrogen | Cat#919788 |
| **MitoTracker Red CMXRos** | Invitrogen | Cat#M-7512 |
| **Wortmannin** | Cayman Chemicals | Cat#10010591 |
| **MitoSOX** | Invitrogen | Cat#M36008 |
| **$H_2O_2$** | Sigma | Cat# 323381 |
| **Critical commercial assays** | | |
| **LDH cytotoxicity kit** | Sigma | Cat#4744926001 |
| **Mouse IL-1 beta/IL-1F2 DuoSet ELISA** | R&D biosystems | Cat#DY401 |
| **Mouse TNF-alpha DuoSet ELISA** | R&D biosystems | Cat#DY410 |
| **Bacterial strains** | | |
| ***Escherichia coli*** | ATCC | 25922 |
| **Software and algorithms** | | |
| **Prism 8.0** | GraphPad | https://www.graphpad.com |
| **BD FACS Diva** | BD Bioscience | https://www.bdbiosciences.com |
| **FlowJo V.10** | BD Bioscience | https://www.flowjo.com/ |
| **FV10-ASW viewer** | Olympus | www.olympus-lifescience.com |
| **Leica LAS AF** | Leica | https://www.leica-microsystems.com/ |
| **FIJI/Image J NIH** | Image J | https://imagej.net/Fiji/Downloads |
| **Experimental Models: Organisms/strains** | | |
| **C57BL/6J (*M. musculus*)** | CSIR CDRI Animal facility | N/A |
| **Mice diet** | | |
| **Chow diet** | Altromin, Germany | Cat#1320 |
| **Plastic ware** | | |
| **48-well flat bottom plate** | Genetix | Cat# 32048 |
| **24-well flat bottom plate** | Genetix | Cat# 32024 |
| **12-well flat bottom plate** | Genetix | Cat# 32012 |
| **6-well flat bottom plate** | Genetix | Cat# 32006 |
| **96-well flat bottom plate** | Genetix | Cat# 32096 |
| **96-well flat clear bottom plate** | Costar | Cat# 3603 |
